# Enhancing pigment production by a chromogenic bacterium (*Exiguobacterium aurantiacum*) using tomato waste extract: A Statistical approach

**DOI:** 10.1101/2024.10.17.618848

**Authors:** Birhanu Zeleke, Diriba Muleta, Hunduma Dinka, Dereje Tsegaye, Jemal Hassen

## Abstract

There is high demand for microbial pigments as promising alternative for synthetic pigments basically for safety and economic reasons. This study aimed at the optimization of yellowish-orange pigment production by Exiguobacterium aurantiacum using agro-waste extract as growth substrate. Air samples were collected using depositional method. Pure cultures of pigment producing bacteria were isolated by subsequent culturing on fresh nutrient agar medium and the potent isolate was identified using MALDI-TOF technique. Culture conditions were screened using Plackett-Burman design and the most three significant variables were optimized by response surface methodology. Fermentation was conducted in 150 mL agro-waste decoctions from which tomato waste extract was selected because of higher optical density of the culture compared to other agro-waste extracts. Pigment was extracted by solvent extraction method and the best solvent was selected based on its ability to dissolve the culture suspension. The pigment was characterized using spectroscopic and chromatographic techniques. Culture agitation rate, initial medium pH and concentration of yeast extract were identified as the most significant (p< 0.0001) variables affecting pigment production. At optimized conditions, 0.96 g/L of pigment was extracted from 4.73 g/L of culture biomass and the extracted pigment under optimized conditions was 1.6 times higher than the pigment extracted under un-optimized conditions. The spectroscopic and chromatographic analyses demonstrated the presence of different functional groups and carotenoids were identified as parts of the molecule responsible for the yellowish-orange pigmentation of the extract. This study demonstrated the potential for optimization of pigment production by bacteria using agro-waste extract as substrate. Hence, the current findings strongly encourage for further study at a large-scale level for industrial production.

## 1. Introduction

The increasing demand for colored products lead to proportional increase of using available synthetic dyes which are not environmentally friendly. Since the middle of 19^th^ century, synthetic dyes replaced natural pigments of higher organisms mainly due to the relatively cheap processing cost and still extensively continued regardless of its deleterious effects [1], [2]. It is estimated that nearly 8 x 10^5^ tons of synthetic dyes are annually produced worldwide and more than 10,000 different dyes are used [3].

Irrespective of the characteristics of the dyes used, the final operations of all dyeing processes involve washing to remove unfixed dyes. The persistent use of synthetic dyes and continual release of dyed effluents to the environment isn’t just a minor inconvenience; rather it’s a major obstacle with profound implications. Because of high stability to light, temperature, water, detergents and chemicals, the effluents persist in the environment for long period of time [4]. The expansion of industries relying on synthetic dyes complemented with huge amount of water consumption that negatively impacts the ecosystem, creating a domino effect that hampers overall productivity and threatens the functions of ecosystems with ultimate negative impact on biodiversity.

Thus, environmental pollution and biodiversity conservation issues demand for eco-friendly natural products from microbes as a substitute of synthetic dyes to fill the gap. Microorganisms produce various pigments through their bioactive compounds and offer promising possibilities for several applications [5]. In view of the possible commercial values of microbial pigments, yield maximization and production cost minimization are the key parameters to be managed to make the production process economically viable [6], [7]. Hence, process optimization is an area of research for the maximum pigment production using cheap agro-wastes as growth substrate. Agro-wastes are composed of cellulose, hemicellulose, lignin, organic acid, salts and minerals which represent the primary source of renewable organic matter as potential substrate for microbial growth [8].

In optimization experiment, the main interest is screening of factors influencing the pigment yield by determining the optimum value of these influential variables [9]. Plackett-Burman Design (PBD) enables screening of process variables for the selection of influential factors from large number of process variables that can be fixed in further optimization processes [10]. PBD has been successfully applied for screening of process components and their effects on production of enzyme by *Bacillus* species from dairy effluent [11], screening of trace nutrients for glycolipopeptide bio-surfactant production [12], screening of process variables in the production of gedunin-loaded liposomes [13], optimization of industrial grade media for improving the biomass production of *Weissella cibaria* [14].

Response surface methodology (RSM) enhances pigment production by reducing the process variables, time and overall cost. It has been successfully applied to optimize the fermentation parameters to investigate the combined effects of process variables by constructing a mathematical model that accurately describes the whole process [15]. RSM was successfully employed to optimize the medium formulation and initial medium pH on lysine–methionine biosynthesis by *Pediococcus pentosaceus* RF-1 [16], enhanced carotenoid pigment production from *Cellulosimicrobium* strain AZ [17], optimization of industrial grade media for improving the biomass production of *Weissella cibaria* [14], assessment of thermal stability of natural pigments produced by *Monascus purpureus* in submerged fermentation [18], optimization of biomass production and cellular growth from psychrotolerant *Paenibacillus* species BPW19 [19] and production of yellowish-orange pigment from *Chryseobacterium artocarpi* CECT 8497 [20].

Thus, the purpose of this research was to screen and optimize culture conditions and media components using PBD and RSM for optimized pigment production by *Exiguobacterium aurantiacum* and evaluating the use of agro-waste extracts as substrate for cultivation. Therefore, in this study, optimum levels of culture conditions and medium concentration that enhanced pigment production were investigated and experimental yield was successfully validated.

## 2. Materials and Methods

### 2.1. Chemicals, Equipment and Reagents

Nutrient agar (NA) medium was used for isolates colony growth and culture cultivation was conducted in liquid agro-waste extracts. Analytical grade organic solvents, obtained from Adama Public Health Research and Referral Laboratory Center, Adama, were used for pigment extraction via centrifugation. The concentrations of dependent variables were measured in absorbance units using photometer 7500, Palintest Wagtech Potalab+ (C), UK and UV-Vis spectrophotometer, VWR P9, China. Aseptic conditions were maintained throughout the experimental activities wherever necessary.

### 2.2. Sample collection, isolation and identification of pigment producing bacteria

The air sample was collected using depositional method according to [21] by exposing nutrient agar plate (Oxoid) to ambient air for 15 minutes. The plate was then incubated at 30°C for 48 hrs. Pure culture of the pigment producing bacteria were isolated by subsequent culturing on fresh nutrient agar medium and the potent pigment producers were screened and identified by MALDI-TOF mass spectrometery technique as per the recommendation of the manufacturers of MALDI-TOF MS systems (Bruker Daltonics, Microflex LT Biotyper operating system) discussed by [22]. Briefly, the colony of pigmented isolate was transferred to Eppendorf tube containing 300 μL of ultrapure water with sterile micropipette tip. Then, 900 μL of absolute ethanol was added and mixed thoroughly. The mixture was then centrifuged at 12000 rpm for 3 minutes and the supernatant was discarded from the Eppendorf tube.

After drying the precipitate at room temperature, 20 μL of 70% formic acid and 20 μL acetonitrile were added and thoroughly mixed. The mixture was centrifuged at 5000 rpm for 3 minutes and the supernatant was transferred to new Eppendorf tube. Finally, after air drying, a microliter of the extracted supernatant was added to the target plate and covered with one microliter of α-Cyano-4-hydroxycinnamic acid (CHCA) matrix solution for analysis.

The characteristic mass spectral fingerprints were generated using EXS3000 MALDI-TOF-MS (Bruker Daltonics, Germany) according to its working manual with laser frequency operating at 60 Hz and, ion source voltage and lens voltage operating at 1.8 kV and 6 kV, respectively. Spectra were generated in the range of 2000 to 20,000 mass-to-charge ratio (m/z). E. coli (ATCC25922) and α-CHCA were used as positive and negative quality control samples, respectively.

For identification, the acquired mass spectra of the isolate was compared with the mass spectra of the known bacteria, the MALDI Biotyper library (MBL), which contains a database of species-specific fingerprints of a wide variety of bacteria to define the taxonomical species to which the isolate belongs. Score values below 1.69 reported as non-reliable genus, scores of 1.70–1.99 were classified as probable genus and scores above 2.00 were probable species [23].

### 2.3. Experimental design for screening and optimization of growth conditions

#### 2.3.1. Culture cultivation on Agro-waste extracts (AWEs)

Growth of the potent pigment producing isolates were examined *in vitro* in locally available twelve AWEs, namely; potato, cabbage, tomato, orange, cannon ball cabbage, onion, watermelon, papaya, carrot, banana and beetroot peels, and bread leftover using growth curve analysis technique [24]. Fruit and vegetable waste extracts are rich in variety of nutritional components such as dietary fibers, proteins and carbohydrates, mineral components like potassium, calcium and magnesium, and vitamins comprised of vitamin A, vitamin C and vitamin E [25], [26], [27], [28].

The agro-wastes were segregated at the point of generation, thoroughly washed, sun dried, grinded into powder and sieved using stainless steel sieve having mesh size of 1 mm to 2 mm. Decoction method was used to release extracts from the agro-wastes with slight modifications according to [29]. Briefly, each grinded agro-waste residues were separately boiled in distilled water at a concentration of 50 g per liter in a working volume of 2 L flasks. Once it reached boiling, the heat was reduced and let simmered for 45 minutes. Finally, the decoctions were strained through gauze fabric and filtered through Whatman filter paper number one, and then transferred into clean stoppered Schott bottles, labelled, autoclaved and stored in refrigerator for successive fermentations as sources of growth substrates for the top selected bacterium.

The potent isolate (coded as PPPI-6) was first inoculated into freshly prepared nutrient broth and incubated over night at 30°C for activation. Fermentation was conducted in a working volume of 150 mL AWEs, each inoculated with 10 mL of one day old culture suspension of the potent isolate at optimized significant culture conditions to evaluate which AWEs best support the growth of the isolate. The culture growth were monitored by measuring the optical density (OD_600_) on daily basis until the growth phase reaches stationary phase. At the stationary phase (after 8 days of cultivation) the OD values of each broth culture were computed and statistically analyzed to identify the nutritious AWEs enhanced more culture growth nutrient broth culture as control. All the tests were performed in triplicates and the values were used as averages of the individual score.

#### 2.3.2. Screening of process variables using PBD

Fermentation was conducted in flasks having capacity of 250 mL with working volume of 150 mL nutrient broth as basal medium to evaluate the impact of each variable and identify the key variables that significantly affect the process. Screening of culture conditions and nutrient concentrations were conducted using Plackett-Burman experimental design, a fractional factorial design used to identify the most significant process variables affecting culture growth [10].

Nine different variables comprising of incubation temperature, initial medium pH, culture agitation rate, incubation period, inoculum age, inoculum size, glucose, yeast extract and salt concentrations were evaluated for their impact on culture growth that was measured in terms of OD_600_ value.

Each variable was represented at two levels, low (-) and high (+) in 12 randomized experimental runs (Table1) as suggested by Plackett-Burman experimental design and incorporated in Design Expert Statistical Software (version 13).

**Table 1.** Independent variables selected for screening using PBD.

| S. No | Variable | Unit | Experimental level |  |
| --- | --- | --- | --- | --- |
|  |  |  | Low (-1) | High (+1) |
| 1 | Incubation temperature | ° C | 23 | 37 |
| 2 | pH |  | 5 | 10 |
| 3 | Culture agitation rate | rpm | 60 | 180 |
| 4 | Incubation period | hr | 24 | 72 |
| 5 | Inoculum size | % | 1 | 2.5 |
| 6 | Inoculum age | hr | 12 | 36 |
| 7 | Yeast extract | % | 0.1 | 1 |
| 8 | Glucose | % | 0.5 | 1.5 |
| 9 | Salt (NaCl) | % | 5 | 10 |

To help reduce the impact of variability and provide a more reliable estimate of the effects of the variables being studied the average value of the response, culture OD values were considered. The main effects model and Pareto analysis were used to identify the most significant factors influencing the response. Based on the magnitude of their effect from ANOVA, the most three significant variables that influence the culture growth were selected for further optimization study.

#### 2.3.3. Optimization of the levels of significant factors using RSM

RSM with face centered central composite design (CCD) was utilized for fitting quadratic model for the response variables to estimate the first-order and second-order terms with three levels matrix (Table 2) by keeping other non-significant variables (incubation temperature, inoculum age, inoculum size, NaCl and glucose) at average values [30], [31]. As maximum pigment production potential correlates with the stationary growth phase, the bacterial culture was incubated until the growth reaches stationary phase which was monitored by measuring the turbidity of the culture at 600 nm [32]. Model assumptions and fitness were checked to evaluate the goodness of fit of the statistical model.

**Table 2.** Design matrix of significant variables for optimization with their corresponding levels.

| Variable name | Units | Low (-1) | High (+1) | -α | +$\alpha$ |
| --- | --- | --- | --- | --- | --- |
| Culture agitation rate ($X_1$) | rpm | 60 | 180 | 60 | 180 |
| pH ($X_2$) | | 5 | 10 | 5 | 10 |
| Yeast extract ($X_3$) | % | 0.1 | 1.0 | 0.1 | 1.0 |

Graphical optimization indicating the interactions between process variables was used to visually analyzing response surface to identify combination of these variables that achieves optimum response value using surface plots [33]. Numerical method using desirability function was applied for simultaneous optimization of all the responses in the process to find the best operating conditions that provide the most desirable response values by setting a goal to find factor settings that maximize the overall desirability [34]. The goals for optimization were set “in range” for the identified input variables to find the best optimal conditions for culture growth and “maximize” for output variables to achieve maximum pigment production.

### 2.4. Verification Test

Prior to optimization, culture cultivation was conducted using the selected tomato waste extract (TWE) as substrate by keeping all the growth conditions at an average value (initial culture pH at 7.5, culture agitation rate at 120 rpm, incubation temperature at 30°C, inoculum age at 24 hrs, inoculum size at 2%, NaCl at 7.5%, glucose at 1% and yeast extract at 0.55%) to identify the effects of the significant variables on bacterial growth and pigment production [30], [31].

For verification, cultivation was conducted in 150 mL of freshly prepared TWE broth at predicted optimal conditions of these significant variables. The bacterial growth kinetics was monitored through OD_600_ measurement by taking 10 mL culture broth with replacement for the withdrawn volume by the same amount of sterile TWE and the graph was plotted against time. At stationary growth phase, the culture was centrifuged using ROTINA 380 R bench top centrifuge (Hettick, Germany) to obtain cell biomass from which crude pigment was extracted. The experiments were conducted in triplicates and the average value of the responses were considered for reliable estimate of the effects of the factors being optimized.

### 2.5. Pigment extraction

Pigment was extracted by solvent extraction method according to [19]. Following centrifugation, the supernatant was discarded and the cell biomass was weighed and re-suspended in variety of organic solvents; methanol, ethanol, acetone and chloroform to choose the best one based on their ability to dissolve pigments in the culture suspension, expressed in pigment intensity from OD measurement values to recover pigment yield.

After re-suspension in respective solvents, the culture suspensions were first vortexed for 10 minutes, heated at 60°C for 30 minutes in water bath and acidified with 3N HCl to enhance extraction [35], [36]. After treatment, the broth culture was centrifuged again at 5000 rpm at 4°C for 20 minutes repeatedly until the residue turned white. Then, residue was discarded and the colored supernatant was filtered through 0.45 µm filter paper and kept in biosafety cabinet (BSC) for a week to evaporate the solvent.

The weight of dry biomass and crude pigment extract were determined using gravimetric technique [37]. Briefly, the biomass was determined by computing change in mass. The crude weight of extracted pigment was determined after evaporating the solvent and further characterized by scanning its absorbance at the wavelength region of 350-750 nm using UV-Vis spectrophotometer [38].

### 2.6. Pigment characterization

The extracted pigment was filtered, concentrated and subjected to different types of spectroscopic and chromatographic analyses for characterization.

#### 2.6.1. Infrared (IR) spectroscopy

Attenuated Total Reflectance-Fourier Transform Infrared (ATR-FTIR) spectroscopic technique was used for spectrum analysis. FTIR spectrometer (Thermo Scientific Nicolet iS50, USA) equipped with ATR device was used to determine the functional groups present in the pigment by passing IR light through the crystal where it is partially absorbed by the sample pressed onto the crystal and then reflects back through the crystal again and travels to the FTIR detector [39].

Two milligram of finely ground solid pigmented extract was placed directly onto the ATR crystal. To ensure good contact between the sample and the crystal, pressure of 700 kg/cm^2^ was applied. The radiation from the IR source of the spectrometer was focused onto the ATR crystal and the output radiation was focused onto deuterated triglycine sulfate (DTGS) crystal detector coated with potassium bromide. The spectrum was collected in the spectral range of 4000–400 cm^−1^ by scanning the sample at an average scan of 32 with resolution of 16 cm^-1^ [40]. Finally, the spectrum was analyzed to determine the sample’s molecular composition.

#### 2.6.2. UV-Vis spectroscopy

The extracted pigment was analyzed using UV-Visible spectroscopy to identify the pigment based on its characteristic absorption peaks. The pigmented extract was dissolved in 99.5% methanol and then 10 mL of the solution was poured into a clean cuvette and inserted into a P9 double beam UV-visible spectrophotometer (VWR, China) which directs a beam of light through the sample using methanol as negative control. After adjusting the absorbance values of the sample between 0.1 and 1.0 absorbance units of the spectrophotometer through dilution, the amount of light absorbed by the sample was measured by scanning the absorbance in the wavelength region of 350-750 nm to find the characteristic absorption peak of the sample which provides valuable information for the identification of pigment type. To ensure consistency and accuracy, measurment readings were taken in triplicates.

#### 2.6.3. Chromatographic analysis

An Agilent 1260 Infinity II LC-MS 6495 (Agilent 1260 Infinity II LC System, Germany) equipped with electrospray ionization (ESI) source and Triple Quadrupole mass analyzer was used to separate the pigment based on its chemical property and identify individual components in the pigment by their mass-to-charge-ratio [41]. Briefly, the sample was first dissolved in 99.5% methanol and centrifuged at 10,000 rpm for 10 minutes at a temperature of 4°C. The clear supernatant was poured into autosampler vial from which 10 μL was injected into the liquid chromatography (LC) system. Chromatographic separation of pigment was conducted using a standard reversed-phase C18 column (10 cm x 4.6 mm, 3µm) set at 35°C before the mass spectra (MS) detection.

After separation, the sample entered the mass spectrometer where it undergoes ionization. Nitrogen gas was used to spray the sample for efficient ionization. The capillary voltage, the gas flow rate, the gas temperature and the nebulizer pressure were set at 3000 V, 5 L/min, 300°C and 50 psi, respectively.

For effective separation and ionization process, 0.1% acetic acid in water and methanol at a flow rate of 1 mL/minute were used as eluents at 40 minutes gradient elution program. The program started with 90% of water with acetic acid and 10% methanol and gradually increased the proportion of methanol from 10% to 90% in the first 35 minutes and remained at 90% methanol for the last 5 minutes before returning to the initial conditions. Spectra were scanned between 50 and 1000 mass-to-charge (m/z) ratio. The generated spectra were compared with database of known compounds to identify possible chromophoric compounds present in the pigmented extract.

### 2.7. Data analysis

Design Expert software (Version 13) was used for variables screening and optimization. All the experiments for AWEs screening were performed in triplicates and data were analyzed using Microsoft Excel spreadsheet. Linear regression analysis was used to estimate the relationship between the independent and dependent variables. One-way ANOVA test was employed to compare the means of the dependent variables across different input variables to see if at least one of the means is significantly different from the others. P-value was calculated by comparing the variance between the groups to the variance within the groups at significance level (α) of < 0.05 as cut-0ff value to conclude that at least one group mean is different from the others [16], [42]. Moreover, post-hoc tests were performed to determine which group means were significantly different from each other using Bonferroni correction method.

## 3. Results and Discussion

### 3.1. Isolation and identification of pigment producing bacteria

Among the colonies grown on NA medium from the air samples incubated at 30°C, an isolate with yellowish-orange pigmented colonies (Fig1 Isolation of pure colony from the mixture of colonies grown on the NA medium for identification) were retrived after 48 hrs of incubation.

**Fig1.**
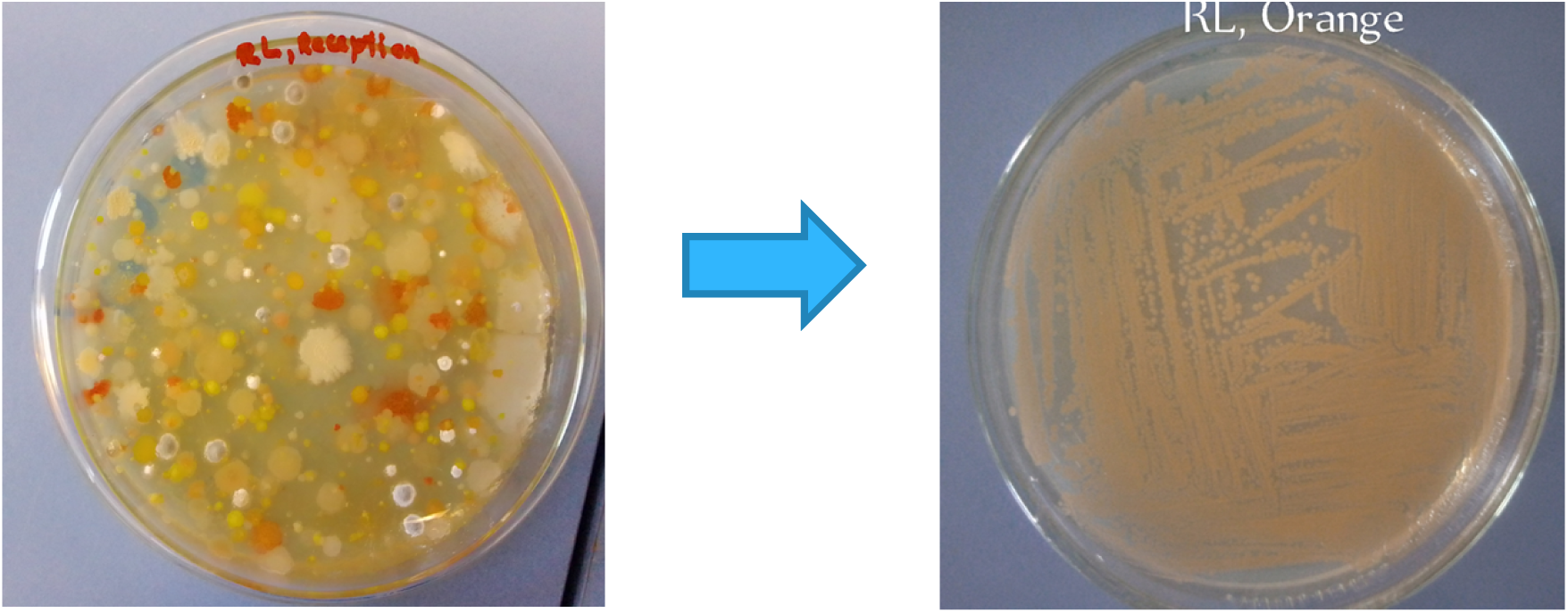

The most potent isolate (PPPI-6) was identified as *Exiguobacterium aurantiacum* by MALDI-TOF mass spectrometry (MS) technique, identified to the species level (Fig 2 MALDI-TOF MS identification result of the potent yellowish-orange pigment producing bacterial isolate to species level as Exiguobacterium aurantiacum) and selected for pigment production through optimization steps using AWEs as growth substrates.

**Fig 2.**
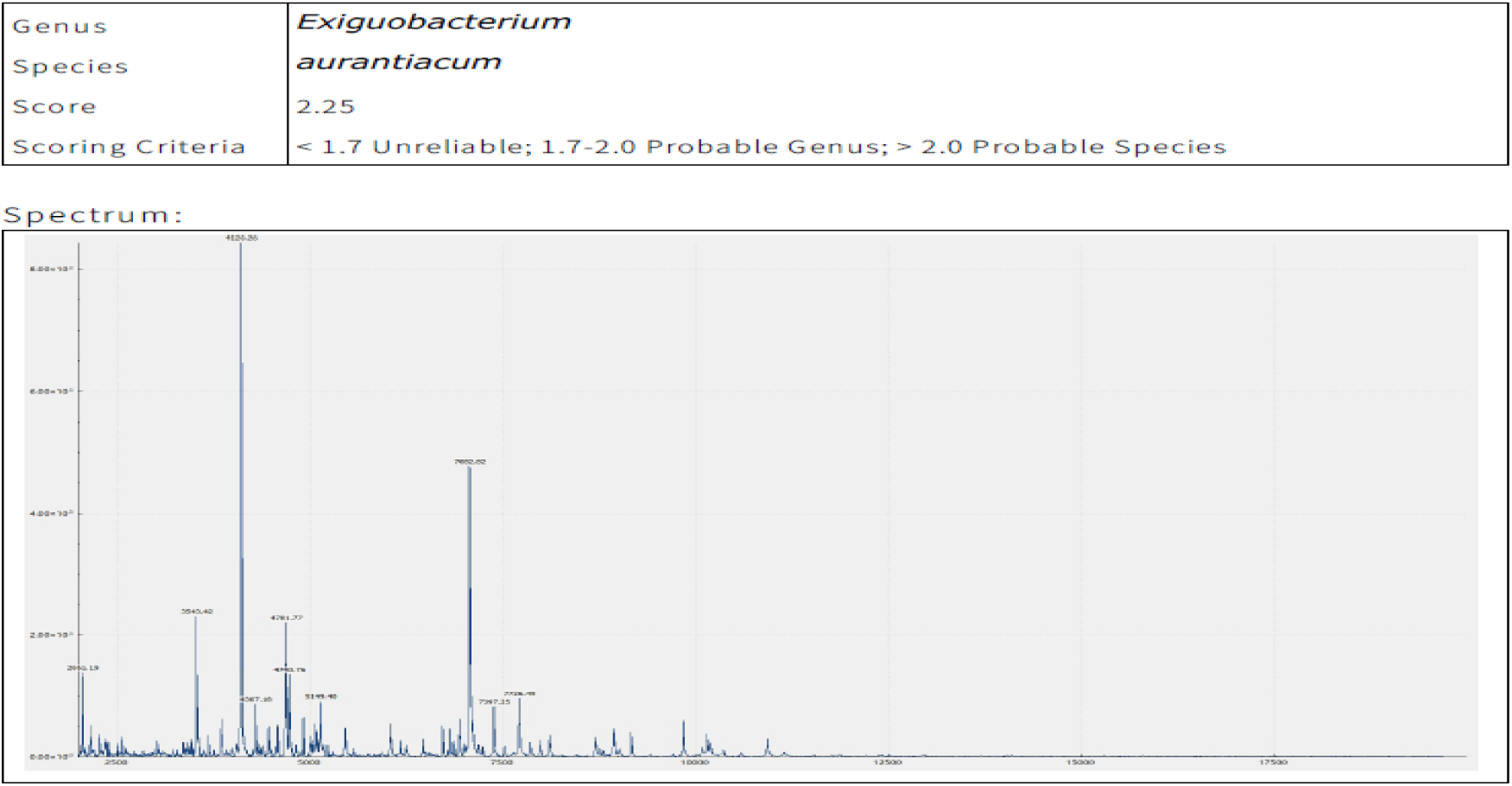

### 3.2. Culture cultivation AWEs

Among the AWEs evaluated for culture growth, three of the agro-waste extracts (banana, beetroot and bread leftover) failed to support the culture growth (OD value less than 0.1) and hence rejected. Accordingly, the rest extracts (Table 3) were utilized as substrates for cultivation using nutrient broth as basal cultivation medium.

**Table3.** Table 3 One-way ANOVA of OD_600_ value of *Exiguobacterium aurantiacum* broth culture.

| Groups | Count | Sum | Average | Variance |
| --- | --- | --- | --- | --- |
| Cabbage | 3 | 1.214 | 0.405 | 3.33E-07 |
| Nutrient broth | 3 | 1.57 | 0.523 | 0.003508 |
| Onion | 3 | 0.641 | 0.214 | 2.33E-06 |
| Papaya | 3 | 0.493 | 0.164 | 1.33E-06 |
| Cannon ball cabbage | 3 | 1.304 | 0.435 | 2.33E-06 |
| Watermelon | 3 | 1.295 | 0.432 | 3.33E-05 |
| Carrot | 3 | 1.004 | 0.335 | 3.33E-07 |
| Tomato | 3 | 1.573 | **0.524** | 4.33E-06 |
| Potato | 3 | 1.385 | 0.462 | 3.33E-05 |
| Orange | 3 | 1.536 | 0.512 | 4E-06 |

ANOVA
| *Source of Variation* | *SS* | *df* | *MS* | *F* | *P-value* | *F crit* |
| --- | --- | --- | --- | --- | --- | --- |
| Between Groups | 0.4313035 | 9 | 0.047923 | 133.49 | 7.19E-16 | 2.392814 |
| Within Groups | 0.00718 | 20 | 0.000359 |  |  |  |
| Total | 0.4384835 | 29 |  |  |  |  |

The one-way ANOVA revealed that there was statistically significant differences in the average OD values between at least two broth cultures of AWEs inoculated with single isolate, F (9, 20 = 133.49, p < 0.05) which indicates there is at least one AWE source which significantly affects the growth of the *Exiguobacterium aurantiacum*. The variation in culture biomass growth under simillar optimized significant process conditions with different AWEs could be attributed to differences in essential nutrient types and composition [43]. Paire wise comparisons of post-hoc test (data not shown) revealed that majority of the extracts have significant difference in OD values. But, tomato versus orange, watermelon versus cannon ball cabbage, watermelon versus cabbage, watermelon versus potato and cannon ball cabbage versus potato waste extracts were not different from each other in the mean culture growth, measured in terms of OD_600_ value.

From OD measurements, tomato waste extract (TWE) showed the highest OD value (Table 3) and therfore, selected as the best low cost cultivation substrate for optimization of pigment production by the chromogenic bacterium (*Exiguobacterium aurantiacum)*. Carbohydrates, proteins, lipids and essential minerals like potassium, magnesium, calcium and iron, and vitamins such as vitamin C, E and various B vitamins are abundant in tomato peel extract [44], [45]. According to [46] tomato peel contains approximately 16.9% soluble sugars, 9.21% cellulose, 10.5% hemicellulose and 42.5% pectin as the most important components ensuring its suitability for microbial growth.

### 3.3. Screening of process variables

A total of nine variables were analyzed with regards to their effects on culture growth (Table 4), expresed in terms of OD value to screen the most significant variables that influence the growth of *Exiguobacterium aurantiacum* using PBD.

**Table 4.**
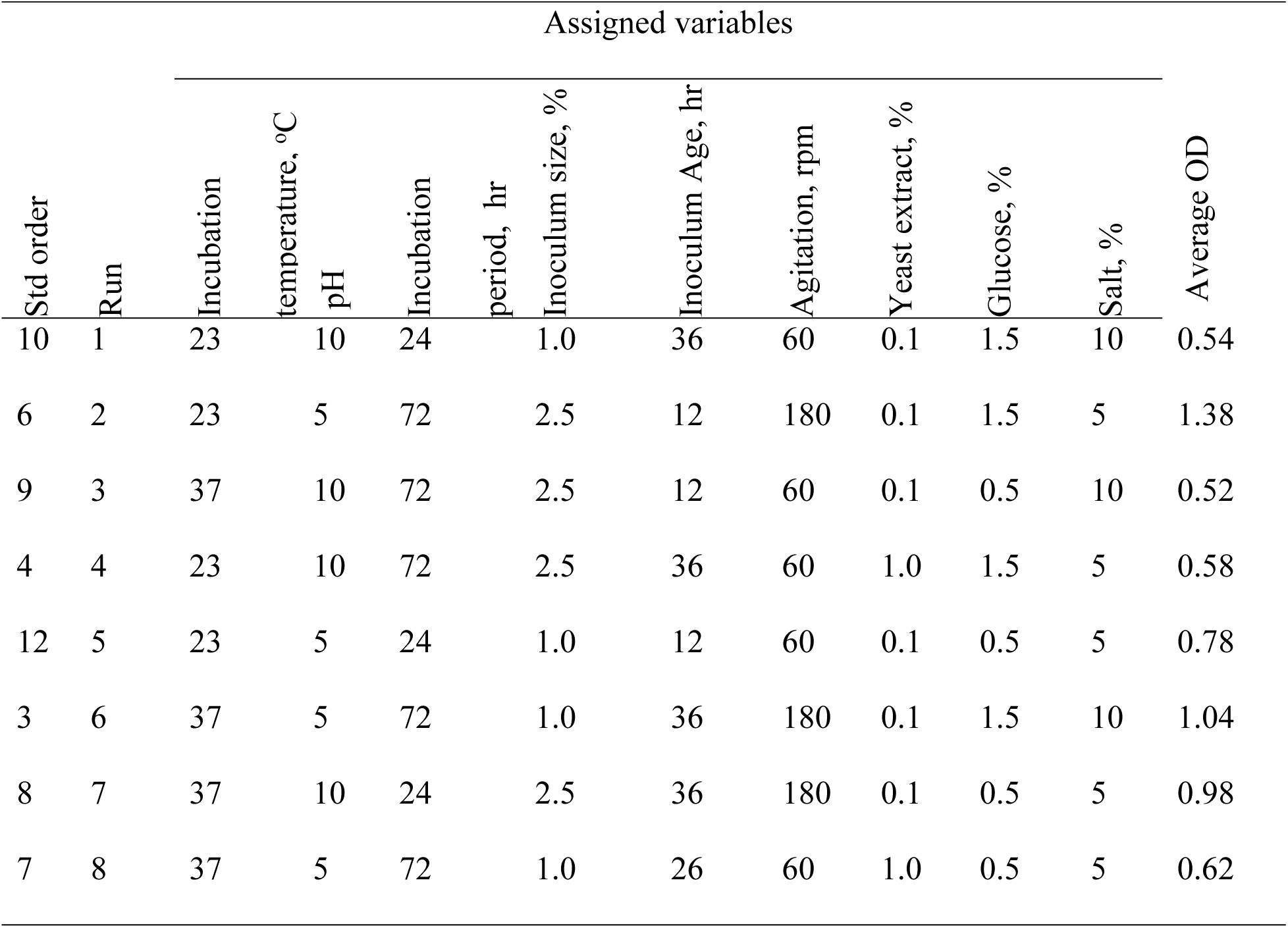
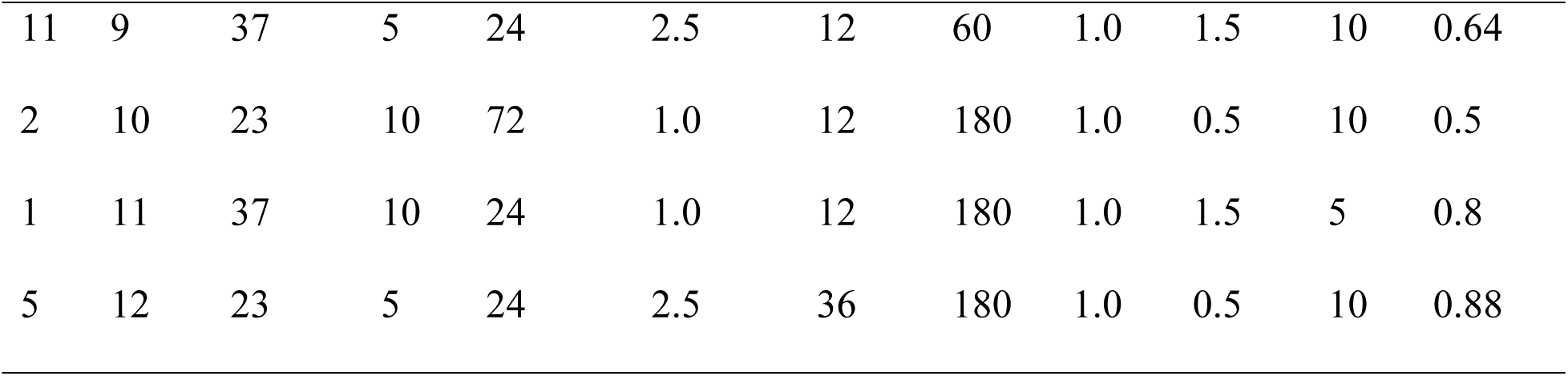
Table 4 PBD experimental run matrix with the corresponding response, OD values Assigned variables.

| Assigned variables |  |  |  |  |  |  |  |  |  |  |  |
| --- | --- | --- | --- | --- | --- | --- | --- | --- | --- | --- | --- |
| Std order | Run | Incubation temperature, °C | pH | Incubation period, hr | Inoculum size, % | Inoculum Age, hr | Agitation, rpm | Yeast extract, % | Glucose, % | Salt, % | Average OD |
| 10 | 1 | 23 | 10 | 24 | 1.0 | 36 | 60 | 0.1 | 1.5 | 10 | 0.54 |
| 6 | 2 | 23 | 5 | 72 | 2.5 | 12 | 180 | 0.1 | 1.5 | 5 | 1.38 |
| 9 | 3 | 37 | 10 | 72 | 2.5 | 12 | 60 | 0.1 | 0.5 | 10 | 0.52 |
| 4 | 4 | 23 | 10 | 72 | 2.5 | 36 | 60 | 1.0 | 1.5 | 5 | 0.58 |
| 12 | 5 | 23 | 5 | 24 | 1.0 | 12 | 60 | 0.1 | 0.5 | 5 | 0.78 |
| 3 | 6 | 37 | 5 | 72 | 1.0 | 36 | 180 | 0.1 | 1.5 | 10 | 1.04 |
| 8 | 7 | 37 | 10 | 24 | 2.5 | 36 | 180 | 0.1 | 0.5 | 5 | 0.98 |
| 7 | 8 | 37 | 5 | 72 | 1.0 | 26 | 60 | 1.0 | 0.5 | 5 | 0.62 |
| 11 | 9 | 37 | 5 | 24 | 2.5 | 12 | 60 | 1.0 | 1.5 | 10 | 0.64 |
| 2 | 10 | 23 | 10 | 72 | 1.0 | 12 | 180 | 1.0 | 0.5 | 10 | 0.5 |
| 1 | 11 | 37 | 10 | 24 | 1.0 | 12 | 180 | 1.0 | 1.5 | 5 | 0.8 |
| 5 | 12 | 23 | 5 | 24 | 2.5 | 36 | 180 | 1.0 | 0.5 | 10 | 0.88 |

From the Half-Normal plot and Pareto chart analysis (Fig 3 Half-normal plot for the standardized effects of process variables on OD value and Fig 4 Pareto chart depicting the influence of process variables on OD value in descending order), respectively, culture agitation rate, initial culture medium pH, yeast extract, salt, glucose and inoculum size were appeared to be significantly larger than the background noise and hence identified as the most significant factors influencing culture growth. The red line (Fig 3 Half-normal plot for the standardized effects of process variables on OD value) fitted to the smallest factors show that the sizes of factors that are likely to be random noise. Whereas, the factors that appeared further to the right of the reference line had larger effects on the response to be included in the mode. Moreover, as indicated in the Pareto chart (Fig 4 Pareto chart depicting the influence of process variables on OD value in descending order), the outlined effects were larger than the red Bonferroni limit and hence those factors were considered statistically significant (p < 0.05, Table 5).

**Fig 3.**
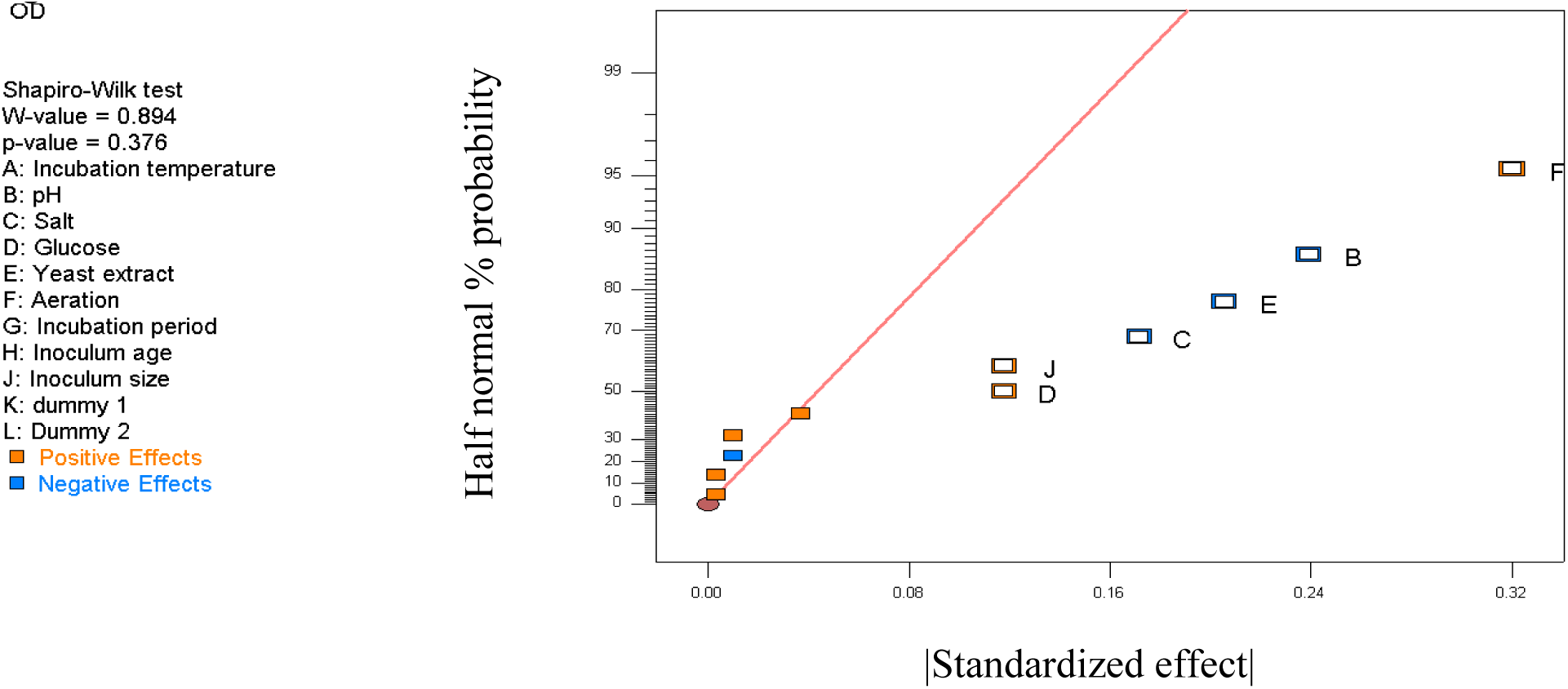

**Fig 4.**
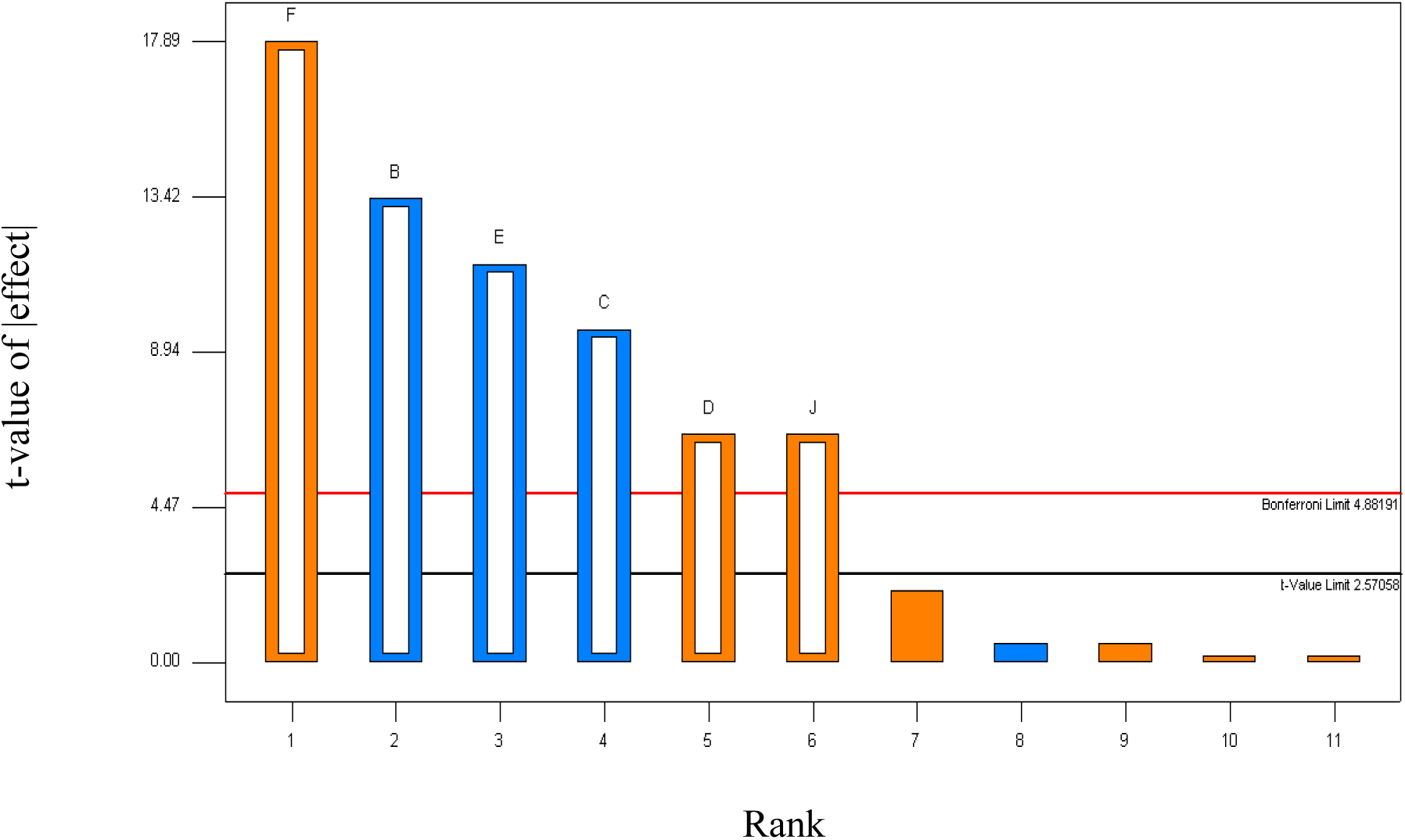

**Table 5.**
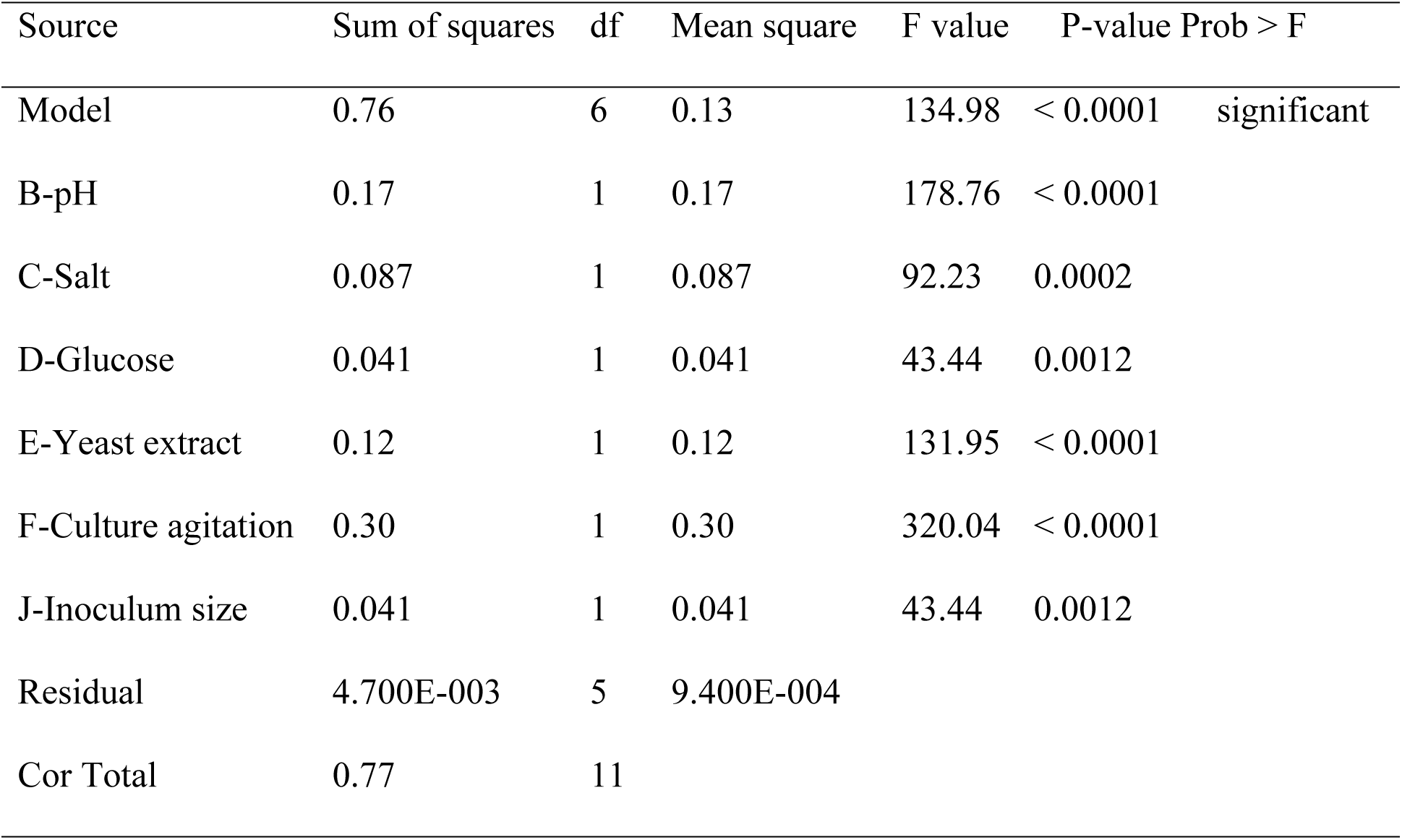
Table 5 ANOVA for selected factorial model.

Experimental data were statistically analyzed by ANOVA test and the model *F* value of 134.98 with the corresponding model P-value < 0.0001 (Table 5) for OD signified that the model was significant. The coefficient of determination (R^2^) value of 99.39% for OD value (Table 6) indicates that up to 99.39% variability in culture growth could be estimated by the selected model terms and the Adjusted R-Squared value of 98.65% is the R-Squared adjusted for the number of model terms. The Predicted R-Squared value of 96.47% also indicates how well future observations may be predicted if the study is repeated under the same conditions and found in good agreement with Adjusted R-Squared show that the model fits well.

**Table 6.** Fit statistics.

|  |  |  |  |
| --- | --- | --- | --- |
| Std. Dev. | 0.031 | R-Squared | 0.9939 |
| Mean | 0.77 | Adj R-Squared | 0.9865 |
| C.V. % | 3.97 | Pred R-Squared | 0.9647 |
| PRESS | 0.027 | Adeq Precision | 36.014 |

From ANOVA data, first-order polynomial model equation was generated using coefficients (Table 7) to predict the response, culture biomass expressed as OD value and written as:

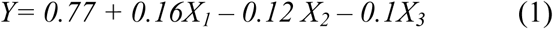

**Table 7.** Coefficients in terms of factors.

| Factor | Coefficient | df | Standard Error | 95% CI, low | 95% CI, high | VIF |
| --- | --- | --- | --- | --- | --- | --- |
|  | Estimate |  |  |  |  |  |
| Intercept | 0.77 | 1 | 8.851E-003 | 0.75 | 0.79 | 1.00 |
| B-pH | - 0.12 | 1 | 8.851E-003 | - 0.14 | - 0.096 | 1.00 |
| C-Salt | - 0.085 | 1 | 8.851E-003 | - 0.11 | - 0.062 | 1.00 |
| D-Glucose | 0.058 | 1 | 8.851E-003 | 0.036 | 0.081 | 1.00 |
| E-Yeast extraction | - 0.10 | 1 | 8.851E-003 | - 0.12 | - 0.079 | 1.00 |
| F-Culture agitation | 0.16 | 1 | 8.851E-003 | 0.14 | 0.18 | 1.00 |
| J-Inoculum size | 0.058 | 1 | 8.851E-003 | 0.036 | 0.081 | 1.00 |

Where Y is OD value, X_1_ is culture agitation rate, X_2_ is initial culture pH and X_3_ is concentration of yeast extract which were selected for further optimization study based on the magnitude of their effects from *F* statistic and p-value of ANOVA test (Table 5).

### 3.4. Optimization of the levels of significant factors

The matrices of each significant variable with their corresponding response values of 20 ace centered CCD experimental run suggested by Design Expert software (Table 8). All the response values; OD (Y_1_), culture biomass (Y_2_) and pigment yield (Y_3_) were recorded at stationary growth phase in set of 20 experimental runs, generated by the software.

**Table 8.** Face-centered CCD experiment design matrices versus their corresponding response values.

| Std | Run | Factors |  |  |  | Responses |  |
| --- | --- | --- | --- | --- | --- | --- | --- |
|  |  | Agitation (rpm) | pH | Yeast extract (%) | OD value | Culture biomass (g/L) | Pigment yield (g/L) |
| 11 | 1 | 60 | 7.5 | 0.55 | 0.65 | 3.6 | 0.82 |
| 13 | 2 | 120 | 7.5 | 0.1 | 0.8 | 4.7 | 0.84 |
| 2 | 3 | 60 | 10 | 0.1 | 0.71 | 4.1 | 0.85 |
| 9 | 4 | 120 | 5 | 0.55 | 0.78 | 4.3 | 0.8 |
| 12 | 5 | 180 | 7.5 | 0.55 | 0.6 | 3.4 | 0.76 |
| 7 | 6 | 180 | 5 | 1 | 0.58 | 3.68 | 0.74 |
| 16 | 7 | 120 | 7.5 | 0.55 | 0.74 | 4.12 | 0.8 |
| 5 | 8 | 60 | 5 | 1 | 0.77 | 4.12 | 0.74 |
| 8 | 9 | 180 | 10 | 1 | 0.89 | 4.2 | 0.88 |
| 1 | 10 | 60 | 5 | 0.1 | 0.88 | 4.62 | 1.06 |
| 19 | 11 | 120 | 7.5 | 0.55 | 0.74 | 4.12 | 0.8 |
| 4 | 12 | 180 | 10 | 0.1 | 0.83 | 4.0 | 0.73 |
| 18 | 13 | 120 | 7.5 | 0.55 | 0.74 | 4.2 | 0.8 |
| 14 | 14 | 120 | 7.5 | 1 | 0.78 | 4.5 | 0.76 |
| 3 | 15 | 180 | 5 | 0.1 | 0.71 | 3.82 | 0.72 |
| 17 | 16 | 120 | 7.5 | 0.55 | 0.74 | 4.2 | 0.82 |
| 20 | 17 | 120 | 7.5 | 0.55 | 0.72 | 4.1 | 0.78 |
| 6 | 18 | 60 | 10 | 1 | 0.78 | 3.92 | 0.65 |
| 10 | 19 | 120 | 10 | 0.55 | 0.84 | 4.2 | 0.78 |
| 15 | 20 | 120 | 7.5 | 0.55 | 0.72 | 4.12 | 0.8 |

The analysis of CCD generated quadratic model for OD (Y_1_) and culture biomass (Y_2_), and two factors interaction (2FI) model for pigment yield (Y_3_) (Table 9) as a function of each variables.

**Table 9.** ANOVA results for responses

| Factor | OD value | Culture biomass (g/L) | Pigment yield (g/L) |
| --- | --- | --- | --- |
| Intercept | 0.7318 | 4.14 | 0.7965 |
| A-pH | 0.0330 | -0.0120 | -0.0170 |
| B-Agitation | -0.0180 | -0.1260 | -0.0290 |
| C-Yeast extract | -0.0130 | -0.0820 | -0.0430 |
| AB | 0.0738 | 0.1775 | 0.0562 |
| AC | 0.0463 | 0.0825 | 0.0312 |
| BC | -0.0037 | 0.0925 | 0.0862 |
| A<sup>2</sup> | 0.0805 | 0.1045 |  |
| B<sup>2</sup> | -0.1045 | -0.6455 |  |
| C<sup>2</sup> | 0.0605 | 0.4545 |  |
| **Model F-value** | 179.14 | 130.58 | 216.10 |
| **Model P-value** | < 0.0001 | < 0.0001 | < 0.0001 |
| **Lack of Fit P-value** | 0.8145 | 0.6927 | 0.9185 |
| **R<sup>2</sup>** | 0.9938 | 0.9916 | 0.9901 |
| **Adjusted R<sup>2</sup>** | 0.9883 | 0.9840 | 0.9855 |
| **Predicted R<sup>2</sup>** | 0.9799 | 0.9678 | 0.9841 |

Coefficients of polynomial equation were computed from ANOVA (Table 9) to predict the values of each response variable and regression equations are presented as:

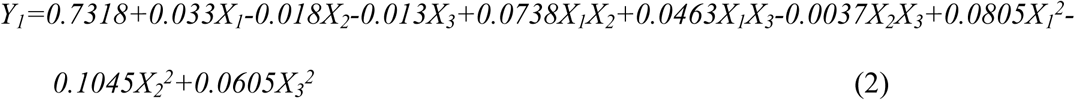

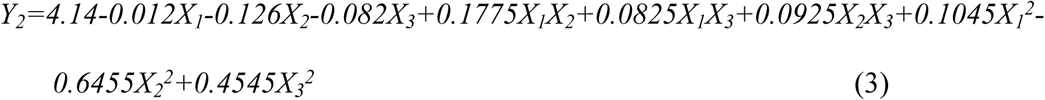

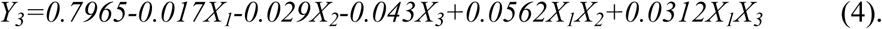

Where X_1_ is culture agitation rate, X_2_ is initial culture pH and X_3_ is concentration of yeast extract.

The ANOVA test revealed that the coefficient of determination (R^2^) (Table 9), representing the proportion of the total variation in the response variable that is explained by the predictor variables in the linear regression model were 0.9938, 0.9916 and 0.9901 for OD value, culture biomass and pigment yield, respectively. The higher the R^2^, the better the model fits the actual data and demonstrates that the influences of the selected variables on response variables are adequately described by the model. Moreover, the larger model *F*-values and smaller model P-value implies the model is significant. The lack of fit p-values were also not significant (p>0.05, Table 9) relative to the pure error that suggests the models were statistically accurate.

### 3.5. Analysis of the effects of the selected culture conditions on response variables

Different ranges of significant factors were used to evaluate their effect on culture growth and pigment production expressed in terms of OD value, culture biomass (g/L) and pigment yield (g/L) (Table 8) and ANOVA test results for response values are summarized in Table 9.

#### 3.5.1. Effects on OD value

The concentration of bacterial cells in nutritious medium (TWE broth) which was estimated by measuring culture OD value was found influenced by culture agitation rate at a linear, quadratic and interaction with initial medium pH (p < 0.0001) (Table 10) but not with concentration of yeast extract (p > 0.05).

**Table 10.** ANOV for Quadratic model. **Response:** OD

| Source | Sum of Squares | df | Mean Square | F-value | p-value |  |
| --- | --- | --- | --- | --- | --- | --- |
| Model | 0.1226 | 9 | 0.0136 | 179.14 | < 0.0001 | significant |
| A-pH | 0.0109 | 1 | 0.0109 | 143.16 | < 0.0001 |  |
| B-Agitation | 0.0032 | 1 | 0.0032 | 42.59 | < 0.0001 |  |
| C-Yeast extract | 0.0017 | 1 | 0.0017 | 22.22 | 0.0008 |  |
| AB | 0.0435 | 1 | 0.0435 | 572.02 | < 0.0001 |  |
| AC | 0.0171 | 1 | 0.0171 | 224.96 | < 0.0001 |  |
| BC | 0.0001 | 1 | 0.0001 | 1.48 | 0.2519 |  |
| A<sup>2</sup> | 0.0178 | 1 | 0.0178 | 234.01 | < 0.0001 |  |
| B<sup>2</sup> | 0.0301 | 1 | 0.0301 | 395.13 | < 0.0001 |  |
| C<sup>2</sup> | 0.0101 | 1 | 0.0101 | 132.13 | < 0.0001 |  |
| Residual | 0.0008 | 10 | 0.0001 |  |  |  |
| Lack of Fit | 0.0002 | 5 | 0.0000 | 0.4263 | 0.8145 | not significant |
| Pure Error | 0.0005 | 5 | 0.0001 |  |  |  |
| Corrected Total | 0.1234 | 19 |  |  |  |  |

Culture agitation enhances oxygen transfer, ensures nutrients are evenly distributed and maintains uniform temperature throughout the culture [47]. Study conducted by [48] have shown that increasing the mixing speed significantly improved bacterial growth rates. Other independent variable which had significant effect on culture growth was linear, quadratic and interaction terms of pH with concentration of yeast extract (p < 0.0001; Table 10). Maintaining optimal pH is crucial to enhance enzymatic activity, achieve best growth and improves fermentation efficiency [49].

The influence due to the interaction of culture agitation and initial culture pH on bacterial culture growth expressed in terms of OD value is illustrated in (Fig 5 Response surface optimization graph (A), Interactive effects between culture agitation (rpm) and pH on OD value). To demonstrate the effects of culture agitation rate, pH and concentration of yeast extract on response variables, response surface graphs were drawn using design expert software to visualize the relationship between each factor and response and type of interactions between test variables.

**Fig 5.**
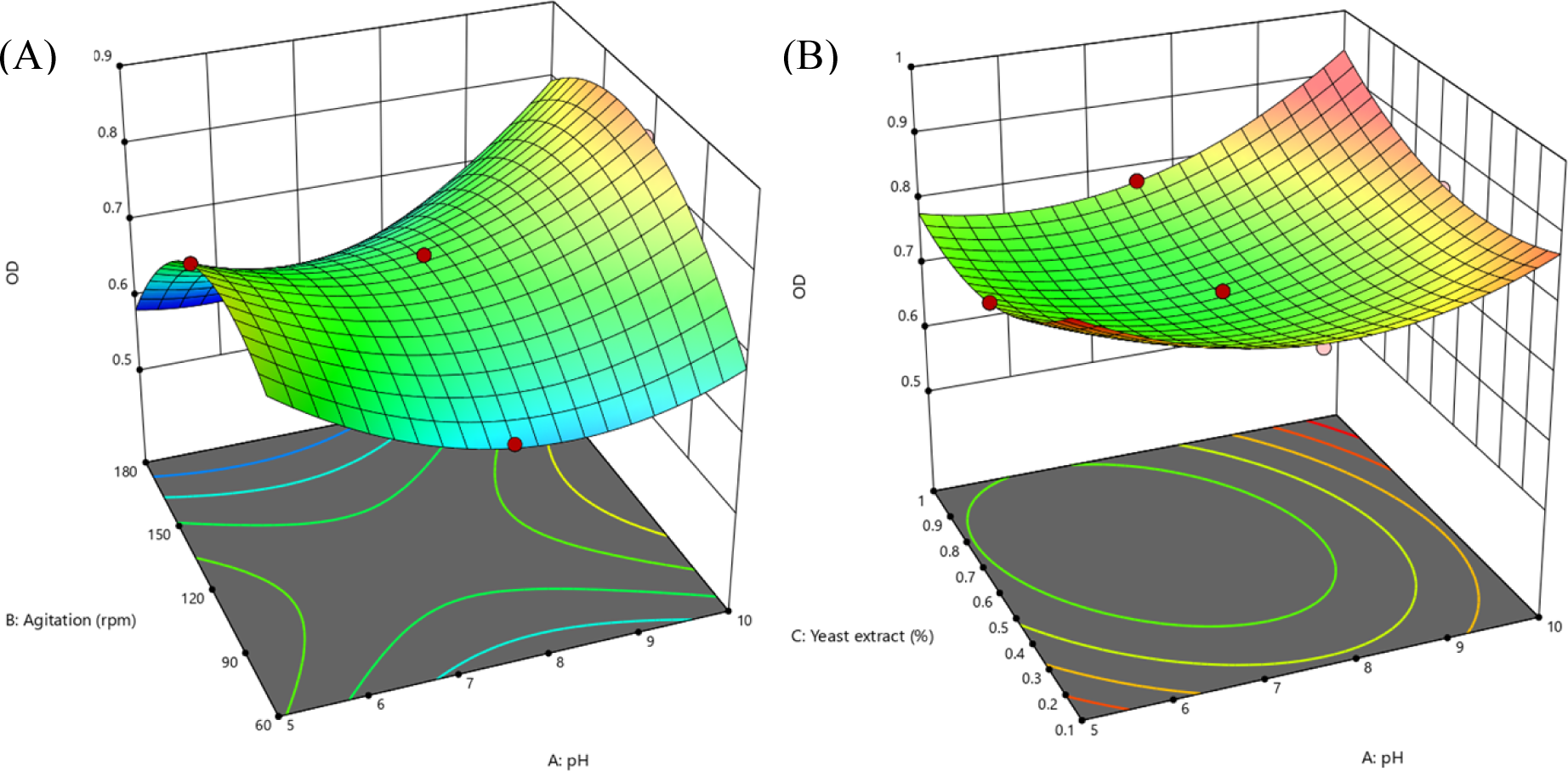
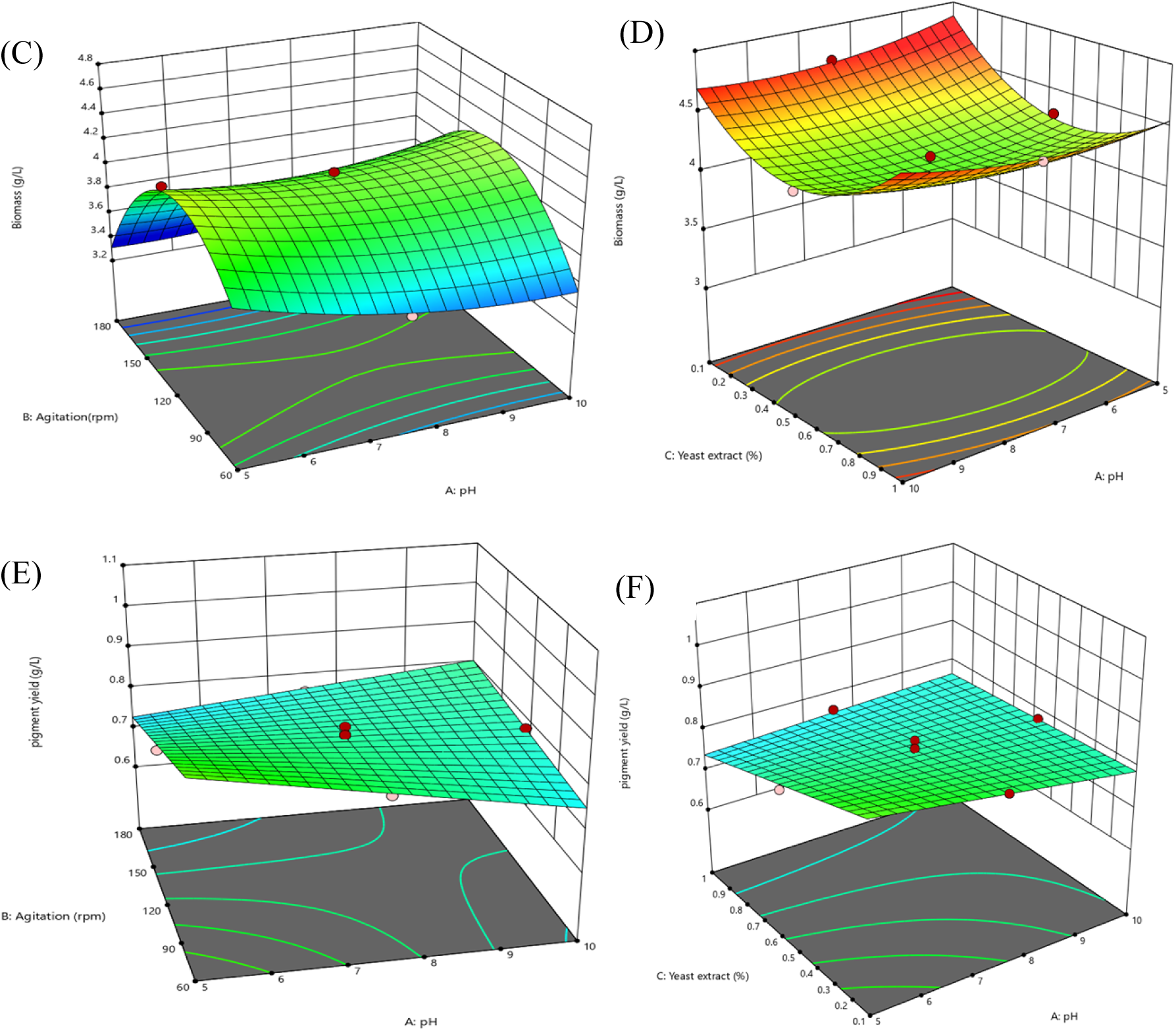

The graphs were generated by varying two independent variables within experimental ranges while keeping the third variable at central point. Fig 5 response surface optimization graphs (Fig 5A, interactive effects between culture agitation (rpm) and pH on OD value, Fig 5C, interactive effects between culture agitation (rpm) and pH on cell biomass (g/L) and Fig 5E, interactive effects between culture agitation (rpm) and pH on pigment yield (g/L)) were generated by varying the culture agitation rate and initial culture pH, keeping the concentration of yeast extract at a central value (0.55%). While Fig 5 response surface optimization graphs (Fig 5B, interactive effects between yeast extract (%) and pH on OD value, Fig 5D, interactive effects between yeast extract (%) and pH on cell biomass (g/L) and Fig 5F, interactive effects between yeast extract (%) and pH on pigment yield (g/L)) were drawn by changing the concentration of yeast extract and initial culture pH, holding the culture agitation rate at a central value (120 rpm).

Maximum OD value of 0.85 was attained at an agitation rate of around 140 rpm and a pH of 10. According to [50], *Exiguobacterim aurantiacum* can grow in a pH range of 5-11 which is in line with the pH range we obtained maximum OD value. With regard to culture agitation rate, our finding also coincides with the recommended agitation rate ranges of 130-150 rpm for bacterial cultures [51].

The combined effects of yeast extract and pH on the OD value is represented by response surface optimization graph, Fig 5B. Both variables exert linear, quadratic and combined effects on OD value of the culture broth. The convex curvature indicates the response surface has a minimum point at which the combination of yeast extract and pH resulted in the lowest OD value. Accordingly, the highest OD values were recorded at the highest values of both yeast extract and pH value followed by the lowest value of yeast extract and both the lowest and highest pH values indicates its pH tolerance. Pertaining to yeast extract concentration in the growth medium, [52] have remarked that even though the optimum concentration can vary depending on specific conditions under which the experiment is conducted, concentration of around 0.7% is effective.

#### 3.5.2. Effects on culture biomass

Relating to culture biomass, culture agitation had a significant effect on culture growth both at linear, quadratic and interaction levels with both pH and yeast extract (Table 11). Other factors which significantly contributed toward culture biomass value were linear and quadratic terms of yeast extract (p < 0.0001), quadratic term of medium pH (p ≤ 0.0015) and combination of yeast extract and pH (p < 0.002) but not linear term of initial culture medium pH (p > 0.05). The growth rate and final biomass of the bacterial culture can impact pigment production and efficient optimization often correlates with increased biomass [53], [54].

**Table 11.**
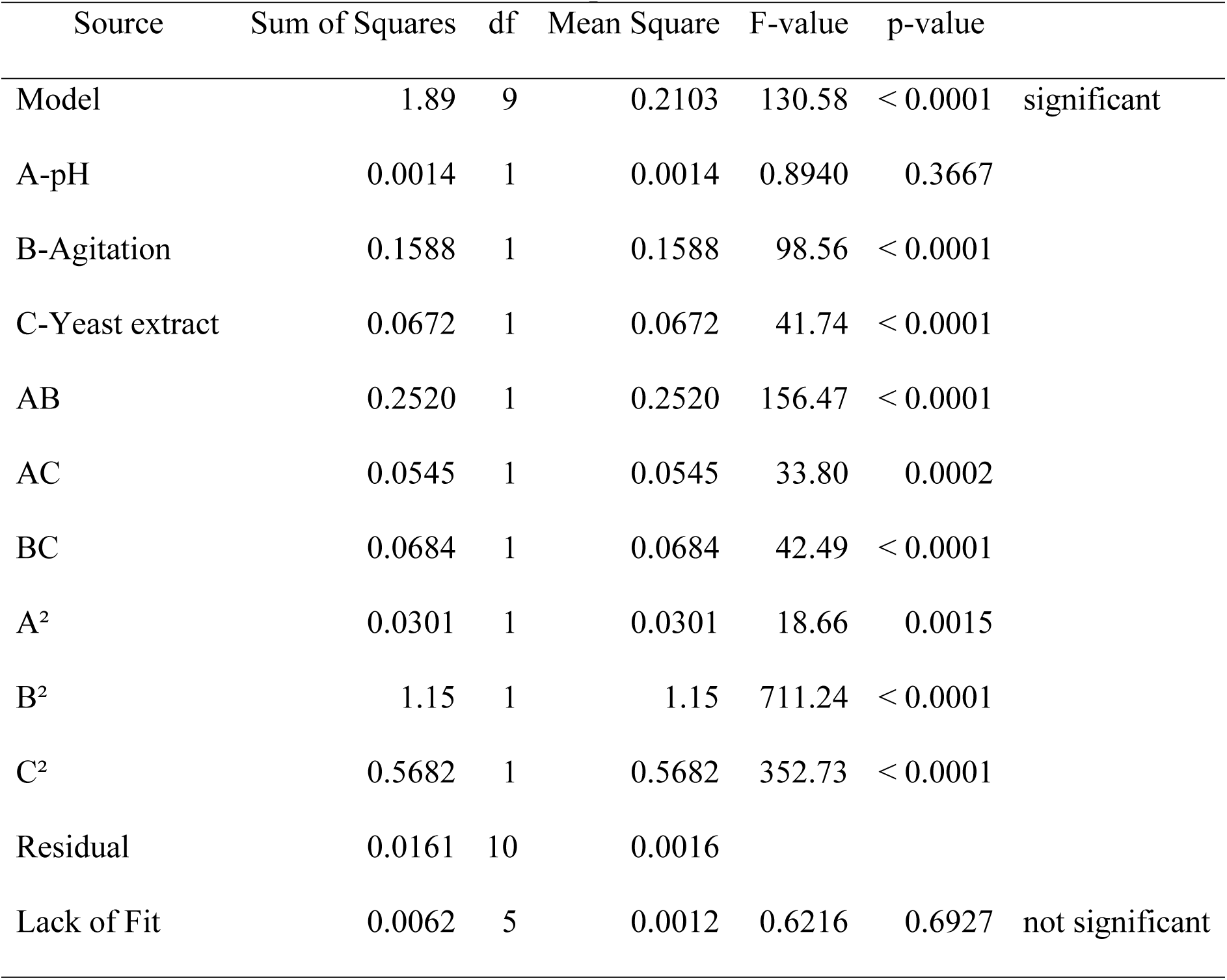
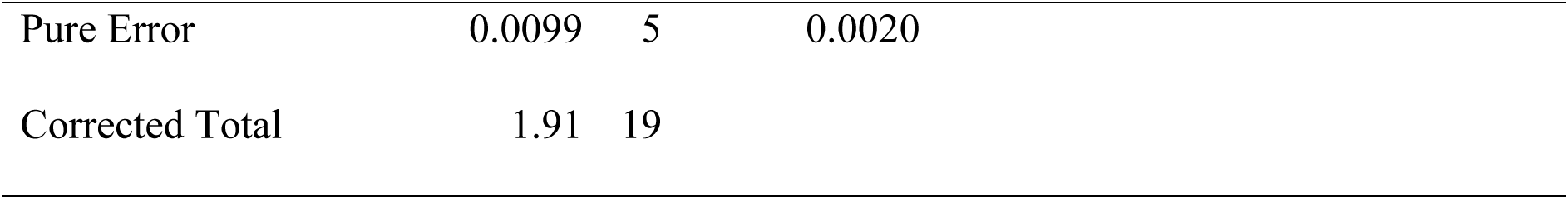
ANOVA for Quadratic model. **Response:** Biomass

| Source | Sum of Squares | df | Mean Square | F-value | p-value |  |
| --- | --- | --- | --- | --- | --- | --- |
| Model | 1.89 | 9 | 0.2103 | 130.58 | < 0.0001 | significant |
| A-pH | 0.0014 | 1 | 0.0014 | 0.8940 | 0.3667 |  |
| B-Agitation | 0.1588 | 1 | 0.1588 | 98.56 | < 0.0001 |  |
| C-Yeast extract | 0.0672 | 1 | 0.0672 | 41.74 | < 0.0001 |  |
| AB | 0.2520 | 1 | 0.2520 | 156.47 | < 0.0001 |  |
| AC | 0.0545 | 1 | 0.0545 | 33.80 | 0.0002 |  |
| BC | 0.0684 | 1 | 0.0684 | 42.49 | < 0.0001 |  |
| A<sup>2</sup> | 0.0301 | 1 | 0.0301 | 18.66 | 0.0015 |  |
| B<sup>2</sup> | 1.15 | 1 | 1.15 | 711.24 | < 0.0001 |  |
| C<sup>2</sup> | 0.5682 | 1 | 0.5682 | 352.73 | < 0.0001 |  |
| Residual | 0.0161 | 10 | 0.0016 |  |  |  |
| Lack of Fit | 0.0062 | 5 | 0.0012 | 0.6216 | 0.6927 | not significant |

|  |  |  |  |
| --- | --- | --- | --- |
| Pure Error | 0.0099 | 5 | 0.0020 |
| Corrected Total | 1.91 | 19 |  |

Provision of rich nutrient sources, maintaining pH within suitable range and proper agitation rates can enhance biomass production [55]. The growth rate and biomass yield of bacterial culture is significantly influenced by variety of nutrient and physical factors and hence adjusting these conditions can optimize culture biomass [49].

The combined effects of agitation versus pH and yeast extract versus pH on culture biomass value are illustrated in response surface optimization graph, Figs 5C and 5D. Culture biomass value increased with the increase in pH while agiration at both lower and higher rates resulted in lower culture biomass yield, response surface optimization graph, Fig 5C. Excessive agitation can cause shear stress, potentially damaging cells and reducing biomass yield [55]. Similarly, lower agitation rate results in lower oxygen transfer rates, creates nutrient gradient and cause cells to clumb together and leads to slower metabolic rate and consecquently lower biomass production [55], [56], [57].

The interactive effect of yeast extract and pH on culture biomass value is shown in response surface optimization graph, Fig 5D. The increasing trend was observed in culture biomass yield at all pH ranges but maximum biomass was recovered at lower yeast extract concentration with slight biomass increament at higher yeast extract concentration. Lower concentaration of nitrogen sources can reduce the risk of nitrogen toxicity which can inhibit microbial growth and promote balanced growth environment that enables the bacteria to efficiently utilize available nutrients for biomass production [52]. *Exiguobacterium aurantiacum* is kown for its adaptability to wide ranges of pH levels which allows the bacteria to maintain metabolic activities and growth across different pH environments by enhancing its overall biomass production [58].

#### 3.5.3. Effects on pigment yield

Pigment yield value was significantly influenced by initial culture pH value, culture agitation rate and concentration of yeast extract at both linear and 2FI levels (p < 0.0001; Table 12).

**Table 12.**
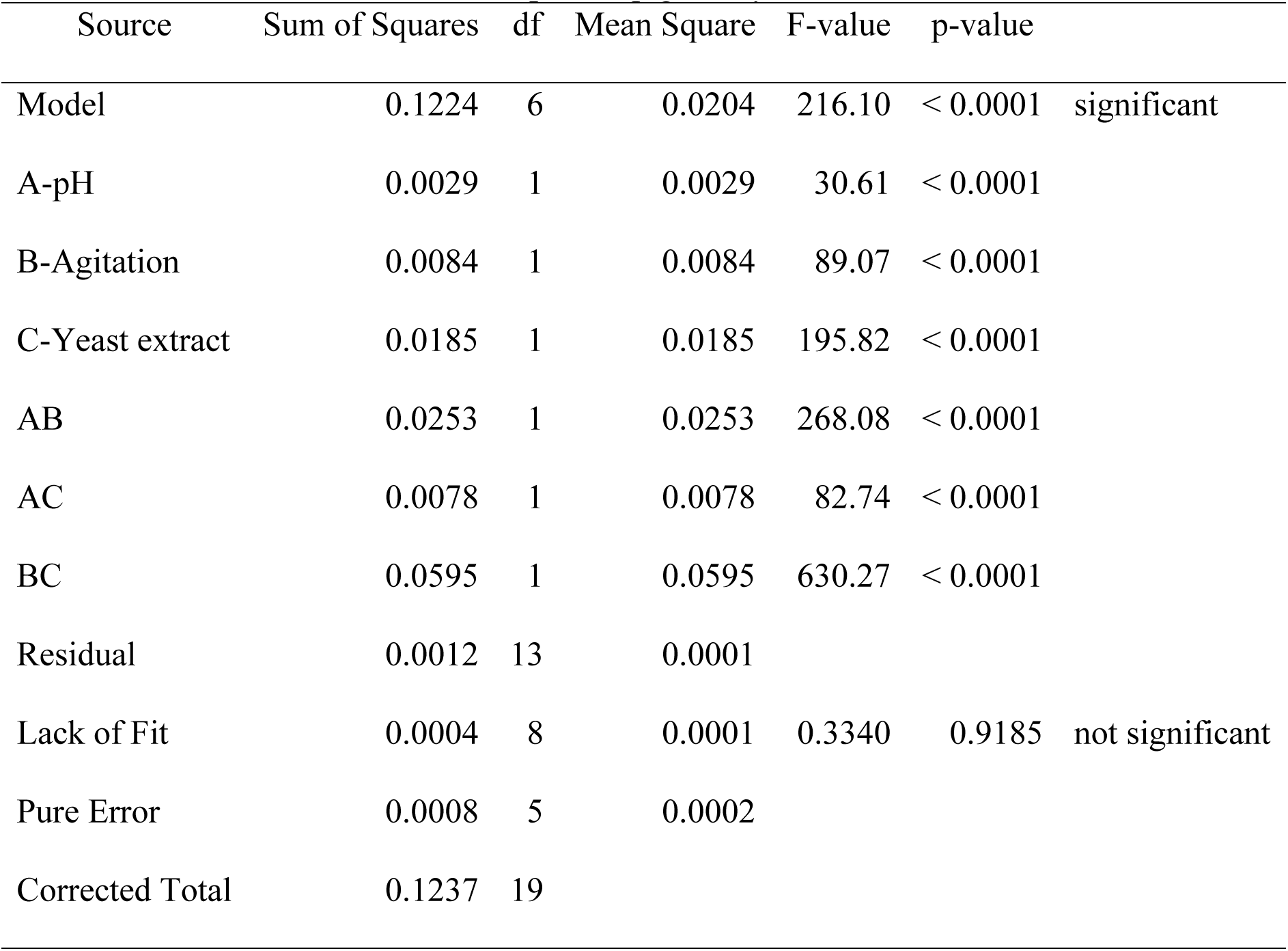
ANOVA for 2FI model. R**esponse:** pigment yield

| Source | Sum of Squares | df | Mean Square | F-value | p-value |  |
| --- | --- | --- | --- | --- | --- | --- |
| Model | 0.1224 | 6 | 0.0204 | 216.10 | < 0.0001 | significant |
| A-pH | 0.0029 | 1 | 0.0029 | 30.61 | < 0.0001 |  |
| B-Agitation | 0.0084 | 1 | 0.0084 | 89.07 | < 0.0001 |  |
| C-Yeast extract | 0.0185 | 1 | 0.0185 | 195.82 | < 0.0001 |  |
| AB | 0.0253 | 1 | 0.0253 | 268.08 | < 0.0001 |  |
| AC | 0.0078 | 1 | 0.0078 | 82.74 | < 0.0001 |  |
| BC | 0.0595 | 1 | 0.0595 | 630.27 | < 0.0001 |  |
| Residual | 0.0012 | 13 | 0.0001 |  |  |  |
| Lack of Fit | 0.0004 | 8 | 0.0001 | 0.3340 | 0.9185 | not significant |
| Pure Error | 0.0008 | 5 | 0.0002 |  |  |  |
| Corrected Total | 0.1237 | 19 |  |  |  |  |

The response surface graph Figs 5E and 5F illustrates the pigment yield as a function of culture agitation versus pH and yeast extract versus pH, respectively. At lower levels of both agitation and yeast extract, pigment yield was significantly increased with decreasing pH which might be attributed to induced stress response at both lower pH and nutrient concentrations, leading to increased production of secondary metabolites and lower agitation can also lead to a more stable environment, reduced shear stress on the cells [36], [59].

### 3.6. Verification tests using TWE as cultivation medium and pigment extraction

Prior to optimization, culture cultivation was conducted in triplicates in 150 mL TWE and 0.65 g/L crude pigment was extracted from culture biomass of 4.20 g/L which had OD value of 0.60 at stationary growth phase taking the average values of each measurment (Table 14).

**Table 13.**
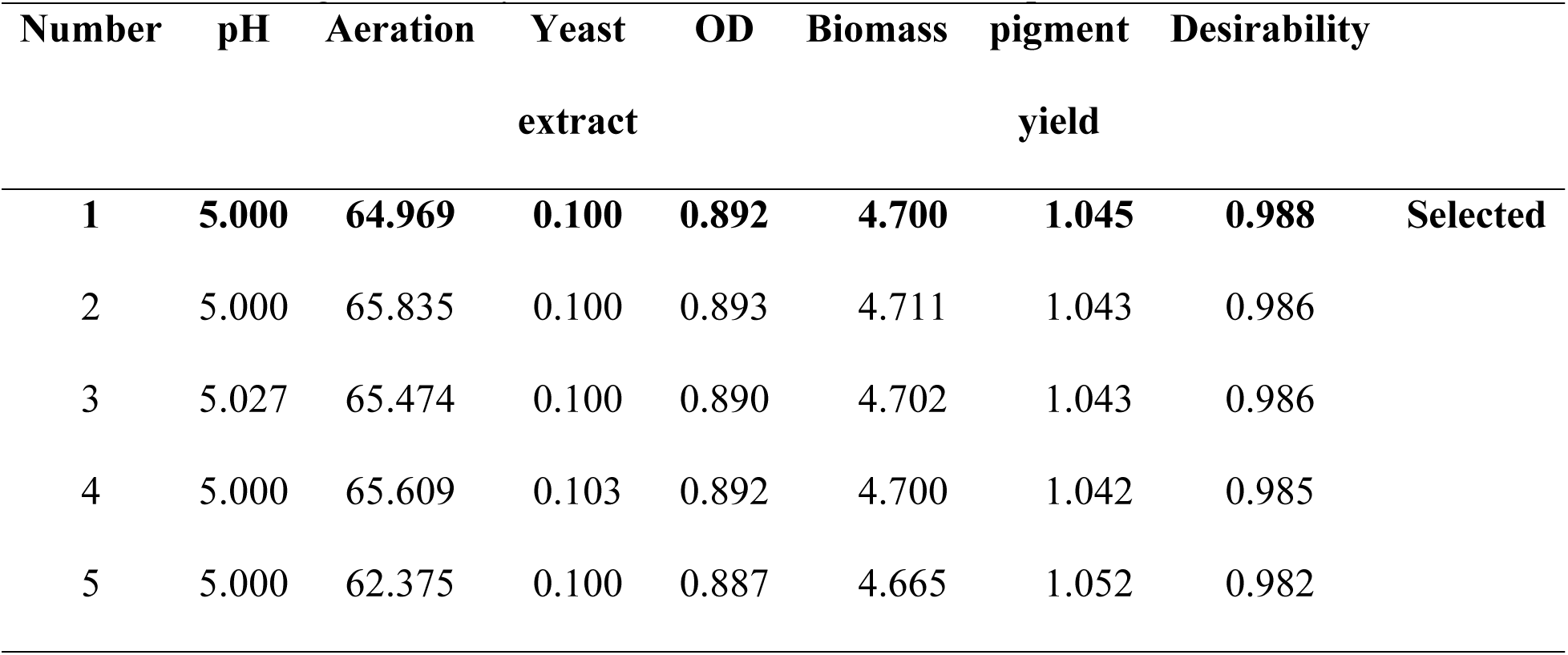
Solutions generated by the software for verification experiment.

**Table 14.** Predicted and experimental values of responses using TWE as substrate at optimized conditions.

| Optimum condition | Coded levels |  | Actual levels |  |
| --- | --- | --- | --- | --- |
| Culture agitation (rpm) | - 0.92 |  | 65 |  |
| Initial culture pH | -1.00 |  | 5.0 |  |
| Yeast extract (%) | -1.00 |  | 0.1 |  |
| Response | Un-optimized value | Predicted value | Optimized value | Optimization efficiency (%) |
| OD value | 0.60 | 0.892 | $0.877 \pm 0.025$ | 31.6 |
| Cell biomass (g/L) | 4.20 | 4.700 | $4.733 \pm 0.076$ | 11.3 |
| Pigment yield (g/L) | 0.65 | 1.045 | $0.957 \pm 0.060$ | 32.1 |

For verification experiment, from the forty-six different solutions (Table 13) generated by the software ranked according to their desirability value, the first solution with pH value equals 5, culture agitation rate equals 65 rpm and yeast extract equals 0.1% at a maximum desirability value of 98.8% were chosen to validate the model.

Culture cultivation was again conducted in the same volume of TWE in triplicates at an optimized significant culture conditions by monitoring bacterial growth dynamics and the broth culture OD_600_ values were measured (Fig 6 Growth kinetics of Exiguobacterium aurantiacum at optimized process conditions using TWE as cultivation substrate) over the incubation period until the broth culture OD value gets stabilized. When the values began to decline, culture biomass was harvested via centrifugation for pigment extraction.

**Fig 6.**
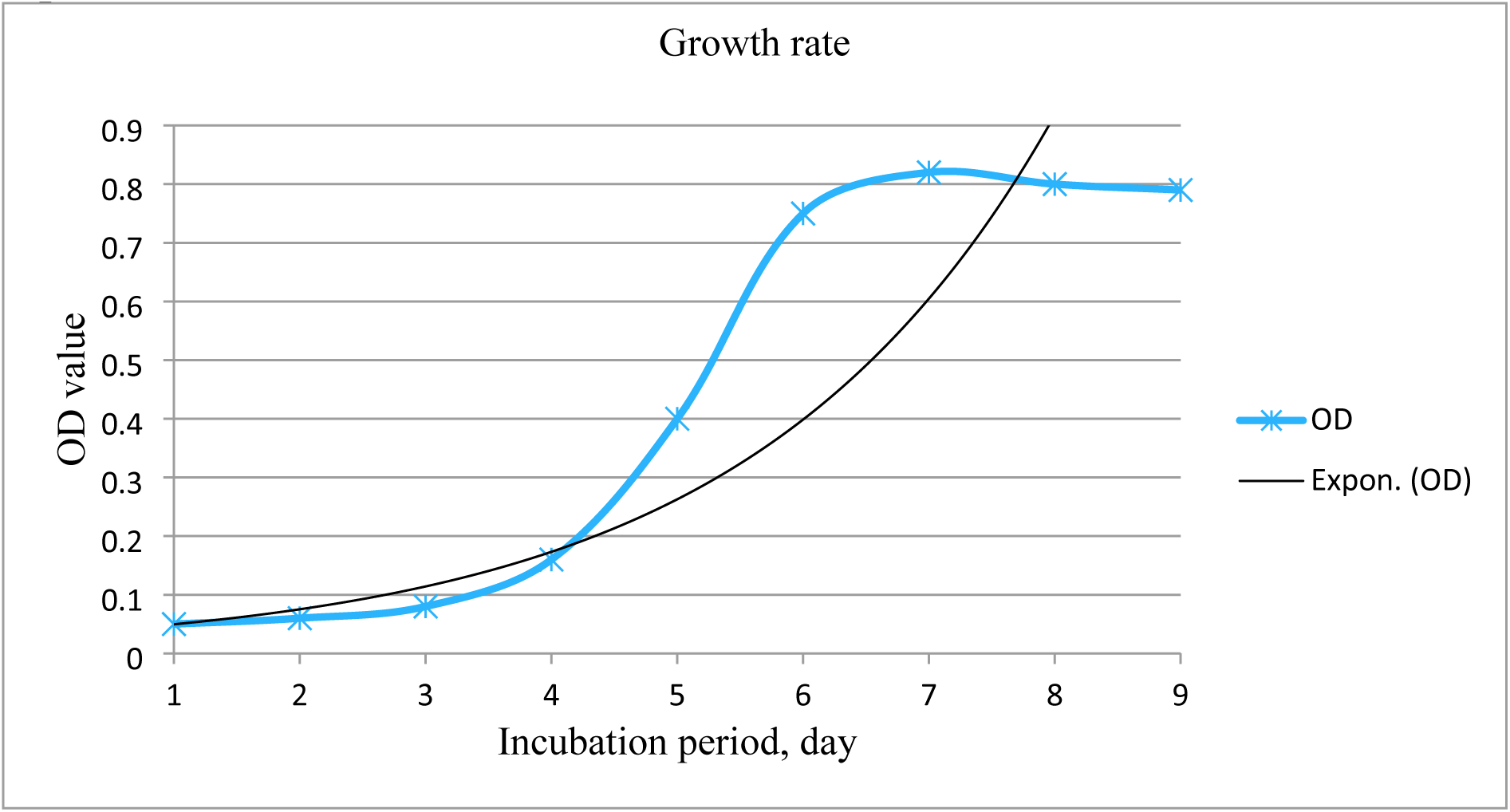

The culture growth was found optimum after 7^th^ day of incubation period when the culture growth reached stationary phase that indicates slow growth which could be attributed to environmental stress [60]. Hence, bacterial biomass was harvested on the 8^th^ day of incubation period in order to recover the intracellular pigment.

The computed biomass and crude pigment yield extracted from the isolate were 4.73 g/L and 0.96 g/L, respectively (Table 14) that revealed the experimental test run produced 0.63% error in biomass concentration prediction which can be attributed to limitation of maintaining exact process conditions and slight amount of noise of the model. Hence, the model was successfully validated as the responses obtained from the actual verification experiment were in good agreement with the results predicted by the model.

Comparing the recovered pigment yield with the result of 180 μg/g biomass obtained from *Chryseobacterium* sp. conducted by [61] after optimization using feather meal as substrate, our yield of 0.96 g/L is better. However, the yield is less than our previous study result extracted from *Micrococcus luteus* using orange waste extract as substrate [62]. The yield variations can be attributed to nutrient compositions of involved culture media and culture conditions of the culture medium used for optimization [63].

### 3.7. Pigment characterization

#### 3.7.1. FTIR spectrum analysis

The IR spectroscopy analysis (Fig 7 The IR Spectrum of pigmented compound extracted from Exiguobacterium aurantiacum.) showed bands of functional groups with peaks at 3270.86, 2921.81, 2852.96, 1738.65 and 1629.69 cm^-1^ and fingerprint regions with different absorption bands, provide insights about the chemical composition of the pigment extracted from the isolate, *Exiguobacterium aurantiacum*.

**Fig 7.**
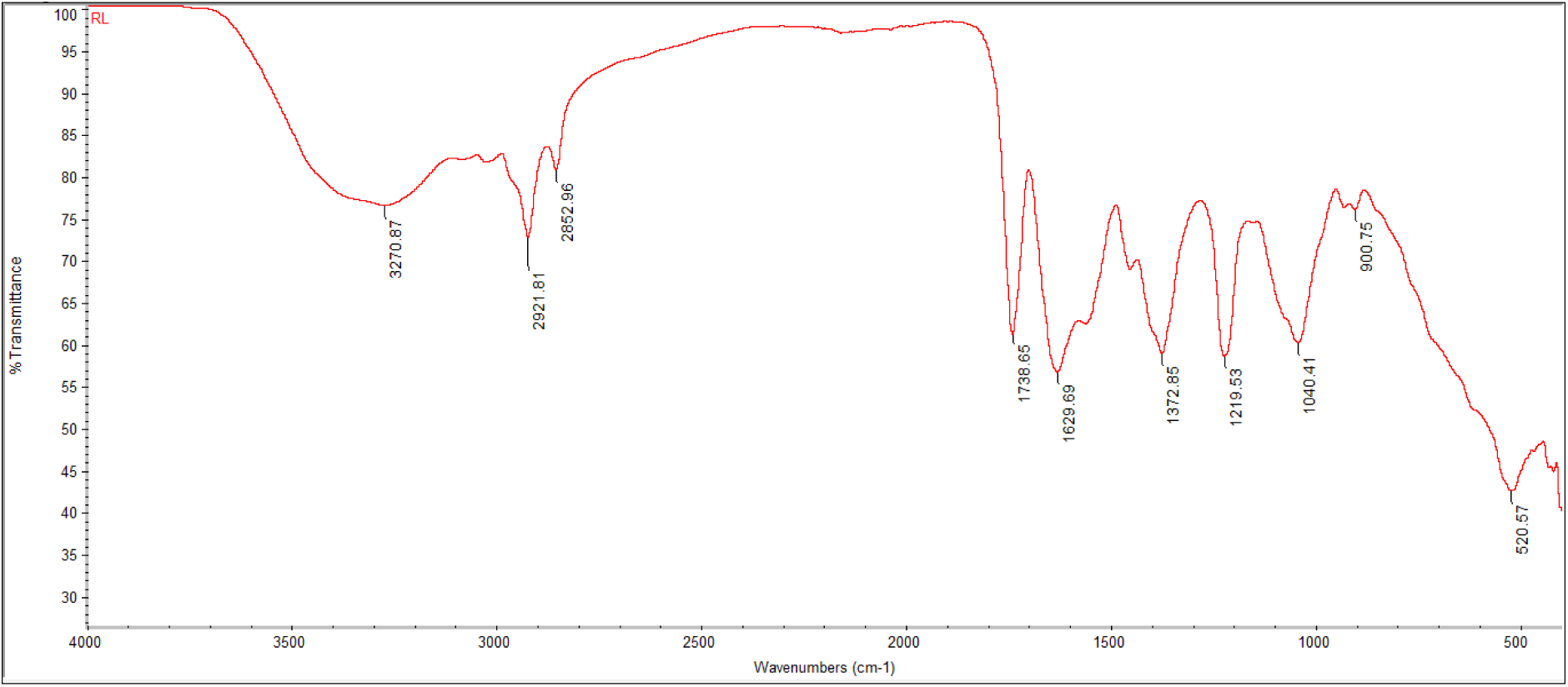

The weak and broad absorption peak at 3270.87.76 cm^-1^ is associated with O-H stretching vibrations, which are characteristic of hydroxyl groups. The second and third short and narrow peaks at 2921.81 and 2852.96 cm^-1^ are indicatives of C-H stretching vibrations commonly found in alkanes. The narrow peak at 1738.65 cm^-1^ is characteristic of C=O stretching vibrations, which are typically found in carbonyl groups and the peak at 1629.69 cm^-1^ is associated with C=C stretching vibrations found in alkenes.

Vibrations help to identify different functional groups in molecules by their characteristic absorption bands [64]. The functional group frequencies analysis revealed that the IR spectrum of yellowish-orange pigment extracted from *Exiguobacterium aurantiacum* indicates the presence of hydroxyl, hydrocarbon, olefinic and carbonyl groups which are among some of the functional groups present in carotenoid compounds [64].

#### 3.7.2. UV-visible spectroscopy analysis

In the present study, methanolic extract of yellowish-orange pigment was subjected to spectral scanning between the wavelengths of 350-750 nm to obtain characteristic absorption peak of the pigment (Fig 8 Absorption spectrum of crude pigment extracted from Exiguobacterium aurantiacum using UV-visible spectrophotometer). Characterizing the absorbance peak of pigment at which it absorbs light most provides information about the chromophores present in the pigment [54], [65].

**Fig 8.**
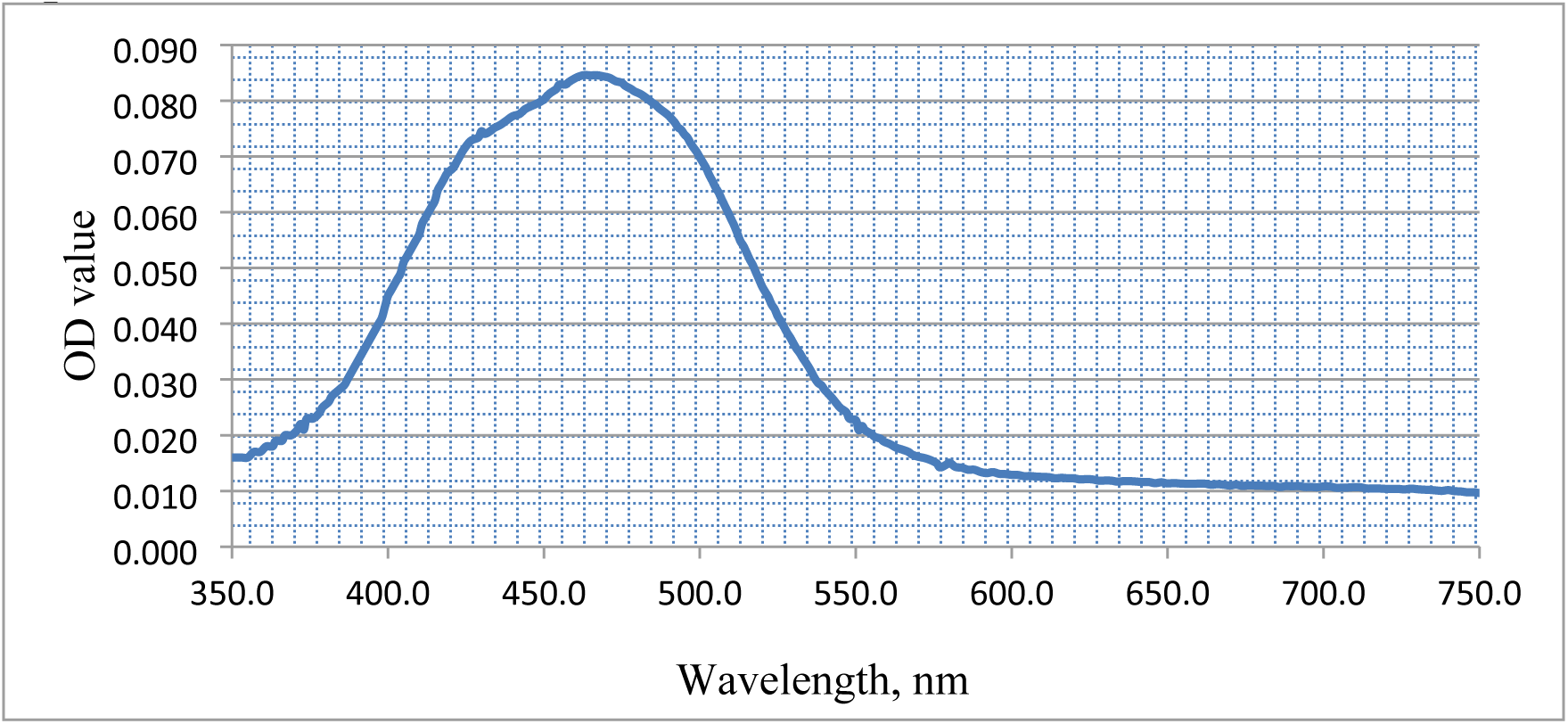

UV-visible spectroscopy result showed that the characteristic absorbance peak of the extract was observed at wavelength of 467 nm that corresponds to the characteristic peak range of carotenoid compounds (400-550 nm), the most important pigment group comprising of yellow to orange-red variants coloration [66], [67]. According to [68] the delocalization of the π electrons of the conjugated system allows carotenoid compounds to absorb specific wavelengths of light in the UV-Visible range of the spectrum.

#### 3.7.3. LC-MS Analysis

The liquid chromatograph (LC) hyphenated with mass spectrometer investigation showed the presence of carotenoid compounds in the analyte, namely; 1,4-Naphthalenedione, 2-(3,7,11,15,19,23,27,31-octamethyl-2,6,10,14,18,22,26,30-dotriacontaoctaenyl)-,(all-E)-and 2,5-Cyclohexadien-1-one,3,3’-(3,7,11,15-tetramethyl-1,3,5,7,9,11,13,15,17-octadecanonaene-1,18-diyl)-bis[6-hydroxy-2,4,4-trimethyl-which is most likely responsible for the vibrant yellowish-orange coloration of the extracted pigment (Figs 9B and 9C, mass spectra of carotenoid compounds extracted from Exiguobacterium aurantiacum, respectively). The total ion current detected over time during the LC-MS run is shown by the plot of Total Ion Chromatogram (TIC) (Fig 9A TIC depicting total ion current detected over time during the LC-MS run), representing the sum of all ion intensities detected at each point in time and the peaks in the TIC correspond to different compounds eluting from the chromatography column at different times. Also, the mass-to-charge ratio (m/z) of ions and their relative abundances are shown by the mass spectra to provide information about the ions detected at that specific time point.

**Fig 9.**
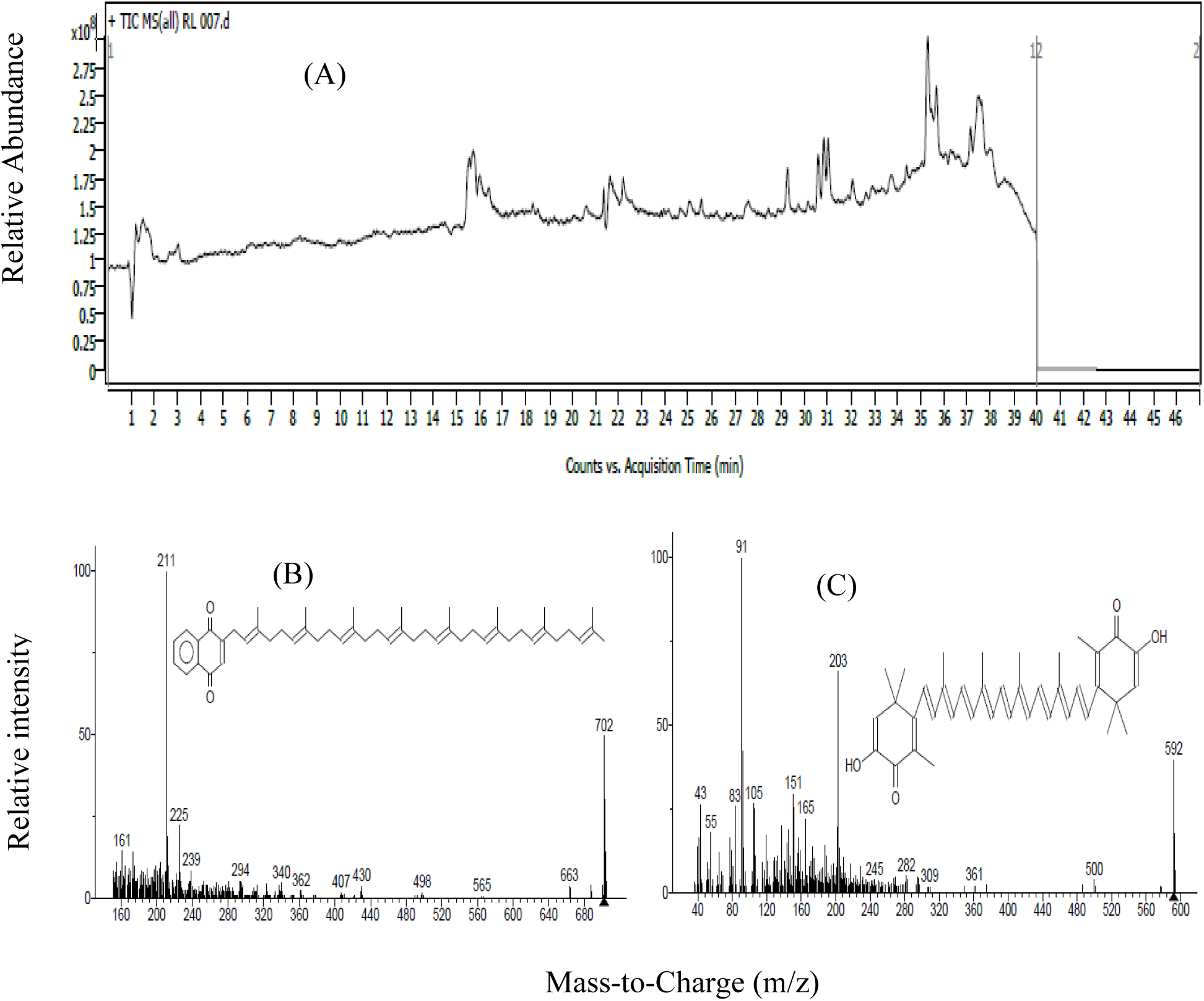

## 4. Conclusion

Eco-friendly natural pigment production and extraction was successfully carried out using TWE as growth substrate. The isolate showed better growth in TWE broth compared to other AWEs used under the optimized significant conditions of initial culture pH, culture agitation rate and yeast extract. The spectroscopic and chromatographic analyses indicated the presence of carotenoid compounds.

## Acknowledgments

The authors are thankful to Adama Science and Technology University, Adama Public Health Research and Referral Laboratory Center, Adama, and Food and Drug Authority and Wudasie Diagnostic Center, Addis Ababa, Ethiopia, for their cooperation to use their laboratory facilities.

## Author Contributions

DM and BZ: Conceptualization and methodology. BZ: Data curation, investigation, original draft preparation, reviewing, editing, formal analysis and visualization. DM, HD and DT: Supervision, validation, project administration, review and editing. JH: Resource, supervision, review and editing. All authors have read and approved the final manuscript.

Funding: This research received no external funding.

Data Availability: Data are included in the article and further clarification can be obtained through the corresponding author if required.

